# Gene Expression Networks in the Drosophila Genetic Reference Panel

**DOI:** 10.1101/816579

**Authors:** Logan J. Everett, Wen Huang, Shanshan Zhou, Mary Anna Carbone, Richard F. Lyman, Gunjan H. Arya, Matthew S. Geisz, Junwu Ma, Fabio Morgante, Genevieve St. Armour, Lavanya Turlapati, Robert R. H. Anholt, Trudy F. C. Mackay

**Affiliations:** Program in Genetics, W. M. Keck Center for Behavioral Biology and Department of Biological Sciences, North Carolina State University, Raleigh NC 27695-7614; Sciome, 2 Davis Drive, Research Triangle Park, NC 27709, USA; Department of Animal Science, Michigan State University, 474 S Shaw Lane, East Lansing, MI 48824; Covance, 100 Perimeter Park, Suite C, Morrisville, NC 27560; Center for Human Genetics and Department of Genetics and Biochemistry, Clemson University, 114 Gregor Mendel Circle, Greenwood, SC 29646; University of North Carolina at Chapel Hill School of Medicine, 321 S Columbia St, Chapel Hill, NC 27516; Key Laboratory for Animal Biotechnology of Jiangxi Province and the Ministry of Agriculture of China, JiangXi Agricultural University, JiangXi, China; Section of Genetic Medicine, Department of Medicine, University of Chicago, Chicago, IL 60637

## Abstract

A major challenge in modern biology is to understand how naturally occurring variation in DNA sequences affects complex organismal traits through networks of intermediate molecular phenotypes. Here, we performed deep RNA sequencing of 200 Drosophila Genetic Reference Panel inbred lines with complete genome sequences, and mapped expression quantitative trait loci for annotated genes, novel transcribed regions (most of which are long noncoding RNAs), transposable elements and microbial species. We identified host variants that affect expression of transposable elements, independent of their copy number, as well as microbiome composition. We constructed sex-specific expression quantitative trait locus regulatory networks. These networks are enriched for novel transcribed regions and target genes in heterochromatin and euchromatic regions of reduced recombination, and genes regulating transposable element expression. This study provides new insights regarding the role of natural genetic variation in regulating gene expression and generates testable hypotheses for future functional analyses.

## Introduction

Understanding how naturally occurring genetic variation affects variation in organismal quantitative traits by modifying underlying molecular networks is a key challenge in modern biology. Most traits are highly polygenic^1–3^ and associated molecular variants have small additive effects on trait variation^4^. Most of these variants are in intergenic regions, up- or down- stream of coding regions, or in introns, and presumably play a regulatory role in modulating gene expression.

Systems genetics analysis seeks to determine how naturally occurring molecular variation gives rise to genetic variation in organismal phenotypes by examining genetic variation in gene expression (expression quantitative trait loci, or eQTLs) and other intermediate molecular phenotypes^2,5–13^. Polymorphic variants associated with variation in gene expression are classified as *cis*- or *trans*-eQTLs depending on whether they are proximal or distal to the gene encoding the transcript, respectively. Genetic variation in gene expression is pervasive; *cis*-eQTLs can have large effects on gene expression that are detectable in small samples; and variants associated with human diseases and quantitative traits tend to be enriched for *cis*- eQTLs^2,5–15^. eQTLs with both *cis-* and *trans*- effects can be assembled into directed transcriptional networks of regulator and target genes^16–18^. Elucidating such regulatory transcriptional networks will facilitate understanding how the effects of individual variants propagate through the network, and how multiple variants together regulate gene expression and affect complex traits^15–18^.

Here, we performed deep RNA sequencing of the *Drosophila melanogaster* Genetic Reference Panel (DGRP) of inbred lines with complete DNA sequences^19,20^. We mapped eQTLs for annotated genes, novel transcribed region (NTRs, which are largely long noncoding RNAs), transposable elements (TEs) and microbiome composition; constructed *de novo cis-trans* eQTL gene expression networks; and evaluated associations of eQTLs and expression traits with organismal phenotypes.

## Results

We collected and sequenced ribo(-) RNA from replicate pools of young flies from each of 200 DGRP lines, separately for males and females. In total we sequenced 1.94 Terabases of RNA, of which on average 13.4 million reads per sample uniquely aligned to the *D. melanogaster* genome (Table S1). The sequences were processed through a pipeline (Figure S1) that (i) removes adapter and rRNA sequences; (ii) aligns and quantifies expressed TE sequences and microbial transcripts; (iii) verifies the origin of each sample; and (iv) quantifies known and novel *D. melanogaster* transcripts and corrects for potential alignment bias due to line-specific sequence variation. We then analyzed normalized expression values for endogenous genes, TEs and microbial species.

### Genetic Variation in Gene Expression

We quantified expression levels of all RNA sequences that aligned to the reference genome in each DGRP line. After elimination of sequences with low expression, we found that 12,806 of 17,097 known *D. melanogaster* genes (75%) were expressed consistently in young adult males and/or females (Table S2A). In addition, we identified 4,282 novel transcribed regions (NTRs) (Table S2B) that showed no overlap with exons on the same strand. A total of 3,846 of the NTRs were located in introns; 290 were anti-sense to known genes, and 146 were intergenic. Most (95.6%) of the NTRs are ≥ 200 bp; the majority (4,149 or 96.9%) lack protein coding potential^21^ (Table S2C) and thus qualify as long noncoding RNAs (lncRNAs)^22–24^. These NTRs in total represent 5.61 Mb new transcribed mature RNA sequences that eluded prior annotation efforts. This increase is likely due to the multiple genetic backgrounds profiled in this study.

Variation in gene expression among the DGRP lines may be confounded by variation in alignment rate to the reference strain due to variation in DNA sequences between the DGRP lines and the reference. Indeed, 2,735 genes (2,117 known genes and 618 NTRs) were affected by alignment bias (Table S2D). We corrected for alignment bias, and partitioned variation in gene expression between males and females, DGRP lines, the sex by line interaction, and residual (environmental) terms (Table S2D), using a false discovery rate of FDR ≤ 0.05. Similar to previous studies^25–27^, we found that gene expression is sexually dimorphic: 98% (96%) of expressed known genes (NTRs) have a significant sex effect (Figure 1A, Table S2D). There is genetic variation in the magnitude of sex dimorphism: 69% (10%) of expressed known genes (NTRs) have a significant sex by line interaction (Table S2D). Therefore, we assessed genetic variation in gene expression separately for males and females (Tables S2D, S2E), and found that 12,151 genes (10,354 known genes and 1,797 NTRs) were genetically variable in females (Figure 1B) and 13,819 genes (11,393 known genes and 2,426 NTRs) were genetically variable in males (Figure 1C). These numbers of genes with significant genetic variation are much higher than previously reported studies, which used microarrays (4,308 in females and 5,814 in males) rather than RNA-Seq^27^. Relative to tiling arrays, RNA-seq has a higher dynamic range and greater precision in quantifying gene expression, although the results from both analyses are positively correlated (Figure S2).

**Figure 1.**
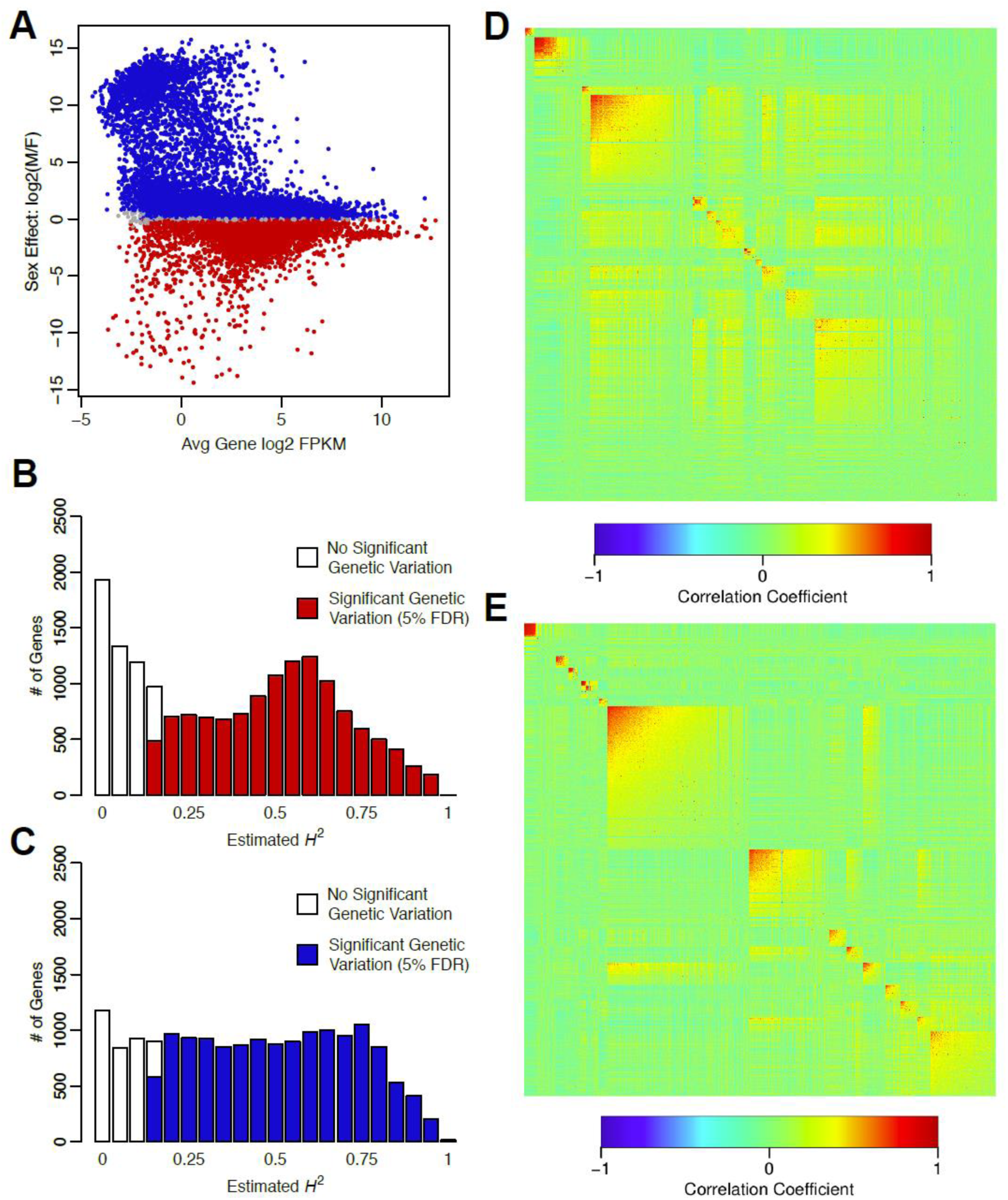
Genetic variation of gene expression in the DGRP. (**A**) Sexual dimorphism of gene expression. Red (blue) indicates significant up-regulation in females (males). (**B**) Distribution of *H*^2^ estimates for annotated genes and NTRs in females. (**C**) Distribution of *H*^2^ estimates for annotated genes and NTRs in males. (**D**) WGCNA modules for annotated genes and NTRs in females. (**E**) WGCNA modules for annotated genes and NTRs in males. Heatmaps show the pairwise correlation of all genes in each module, sorted by average connectivity, with the most tightly connected module at the top left.

Broad sense heritabilities (proportion of phenotypic variance due to genotype differences) ranged from *H*^2^ = 0.148 – 0.986 in females and *H*^2^ = 0.145 – 0.986 in males (Figures 1B, 1C). Notably, 472 (514) of the genetically variable genes in females (number of for males in parenthesis) were located in molecularly defined heterochromatin (*2LHet*, *2RHet*, *3LHet*, *3RHet*, *XHet*, and *YHet*) and chromosome *4*. While there are 6.92× (5.52×) as many annotated genes relative to NTRs in euchromatic regions in females (males); there are 2.21 × (3.18×) as many NTRs in heterochromatin and chromosome *4* in females (males) (Table S2G). Thus, NTRs are highly enriched in heterochromatic regions.

We used weighted gene co-expression network analysis (WGCNA)^28^ to assess the extent to which gene expression levels are genetically correlated in each sex (Figures 1D, 1E, Table S3). We found 13 (15) co-expression modules in females (males). We assessed the extent to which each module was significantly enriched^29^ (FDR ≤ 0.05) for gene ontology (GO) terms and pathway and protein domain annotations (Table S3). For example, female Module 2 (149 genes) is enriched for GO terms involved in ovary function and male Module 6 (365 genes) is enriched for biological process GO terms involved in male reproduction. Female Module 12 (88 genes) and male Modules 13 (35 genes) and 14 (165 genes) are enriched for GO terms affecting small molecule metabolism. Female Modules 3 (26 genes), 6 (27 genes), and 7 (21 genes) and male Modules 9 (42 genes) and 12 (44 genes) are enriched for GO terms affecting innate immunity, and female Module 13 (560 genes) is enriched for GO terms affecting chemosensation.

### Gene Expression QTLs (eQTLs)

We performed genome wide association eQTL analyses for each of the genetically variable genes in each sex. We used ∼1,932,427 million common (minor allele frequency > 0.05) polymorphisms and accounted for effects of Wolbachia infection, polymorphic inversions and polygenic relatedness on gene expression^20,27^. We mapped 90,634 eQTLs in females and 147,412 eQTLs in males (FDR ≤ 0.05). A total of 2,053 genes in females (1,818 known genes and 235 NTRs) and 3,178 genes in males (2,790 known genes and 388 NTRs) were associated with at least one significant eQTL. We defined potentially *cis*- and *trans*-regulatory eQTLs as ≤ 1 kb and > 1 kb of their respective gene bodies. We mapped *cis*-eQTLs to 1,435 (2,071) genes in females (males) (Tables S4A, S4B) and *trans*-eQTLs to 1,527 (2,281) genes in females (males).

We visualized the significant eQTLs by plotting the polymorphism positions on the *X*- axis and the gene positions on the *Y*-axis such that the diagonal corresponds to *cis*-eQTLs and the off-diagonal to *trans*-eQTLs (Figure 2A). The eQTLs tend to cluster in LD blocks in pericentromeric regions, where recombination is suppressed (Table S4C). Many *cis*-eQTLs are also *trans*-eQTLs as indicated by vertical off-diagonal *trans*-bands. We also observed genes associated with many eQTLs throughout the genome, visualized as horizontal off-diagonal *trans*-bands (Figure 2A). In females (males), 217 (377) genes have 200 or more eQTLs, and 22 (43) genes each have greater than 1,000 eQTLs (Tables S4D, S4E). Genes with 200 or more eQTLs are more likely to be found in heterochromatin than those with fewer than 200 eQTLs, and are more likely to be NTRs than annotated genes (Figure 2B, Table S4F). Genes with 200 or more eQTLs that are located in euchromatin are more likely to be located in pericentromeric regions at the border of heterochromatin where recombination is reduced^30^ than those with fewer eQTLs; they are also more likely to be NTRs (Figure 2A, Table S4F).

**Figure 2.**
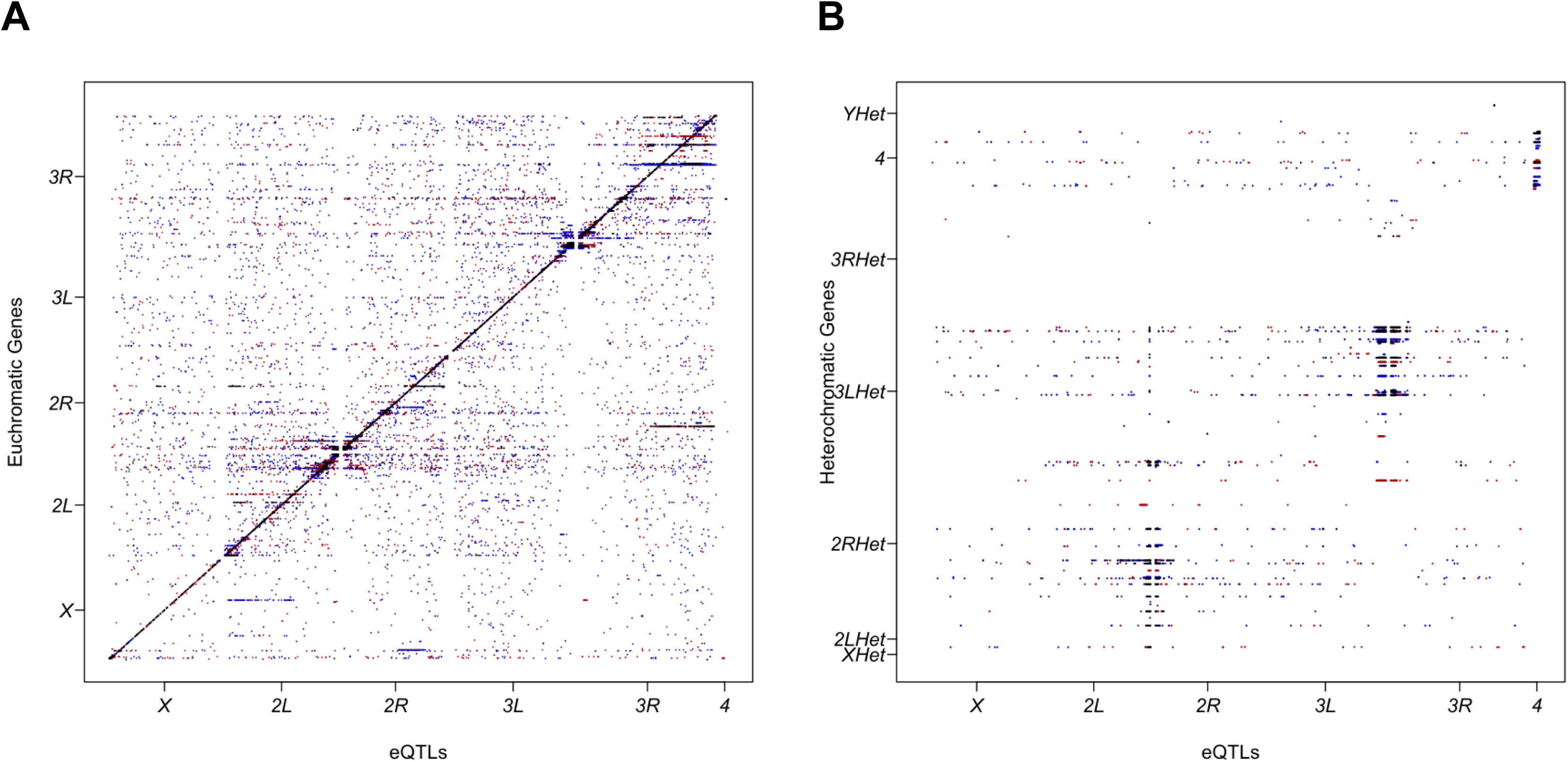
Genomic location of eQTLs for gene expression and genes they regulate. eQTL chromosome positions (bp) are given on the *X*-axis, and the genes with which they are associated on the *Y*-axis. Red points denote female-specific eQTLs, blue indicates male-specific eQTLs, and black shows eQTLs shared by males and females. (**A**) Euchromatic genes. (**B**) Heterochromatic genes.

### eQTL Regulatory Networks

The existence of eQTLs that are *cis*-eQTL for gene X and also *trans*-eQTL for gene Y (Tables S5A, S5B) enables us to construct gene regulatory networks based on multifactorial variation in a natural population. We identified 408 (794) such regulatory interactions supported by at least one *cis*-*trans* eQTL connecting 257 (471) regulatory genes (*cis* end) to 251 (447) target genes (*trans* end) in females (males) (Tables S5C, S5D). There are two or three large regulatory networks in each sex, and many smaller networks (Figures S4, S5). The regulatory genes are largely distinct between the two sexes, although many target genes are in common between males and females (Figures 3, S3, Table S5E). Genes from the sex-specific regulatory networks or from the common networks are not enriched for any GO terms. It is not clear from their anatomical gene expression patterns how the sex-specificity could arise, since the majority of these genes are expressed in multiple tissues, including the reproductive tissues of both sexes^31^.

**Figure 3.**
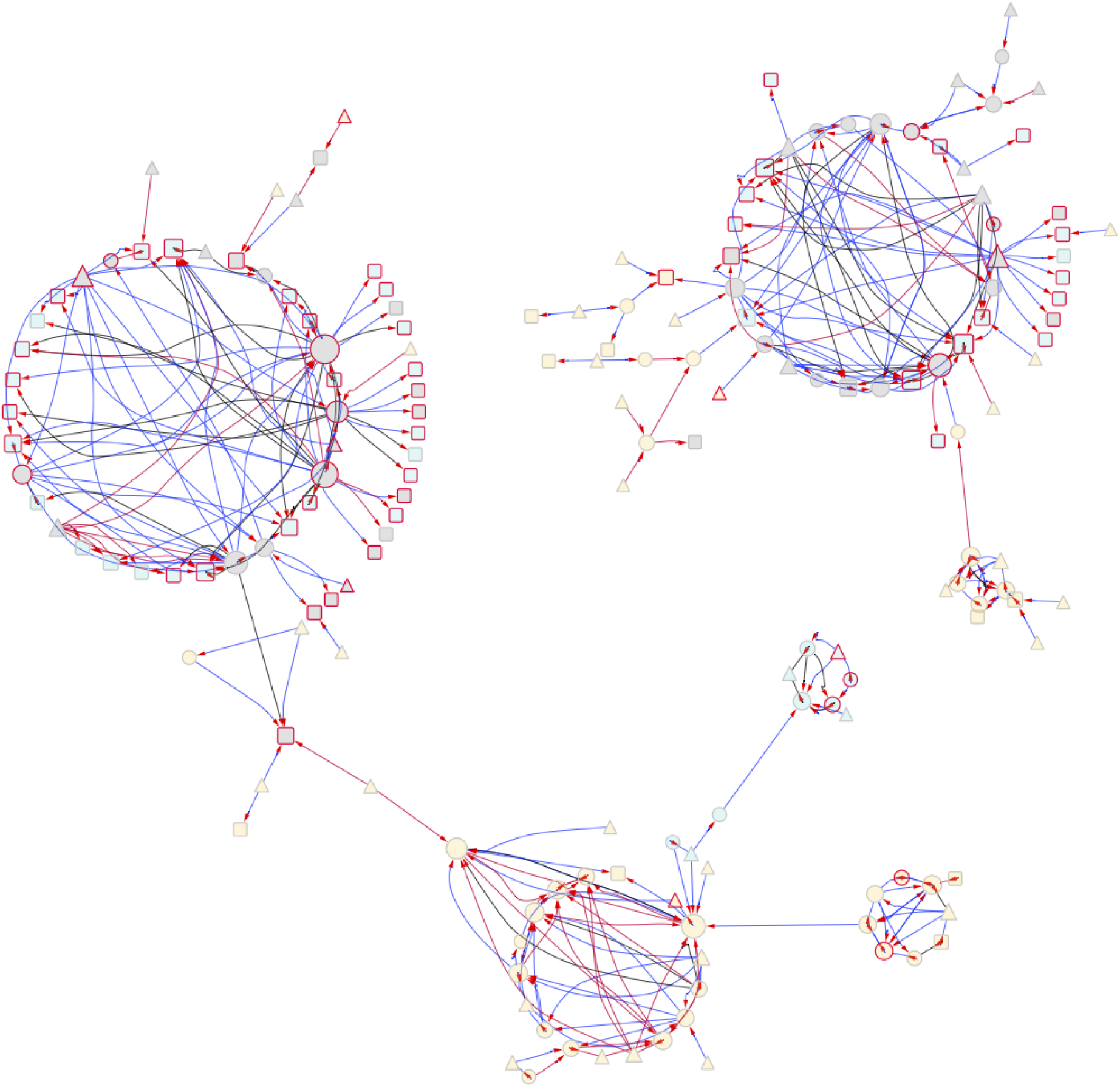
Large *cis*-*trans* eQTL genetic network in females and males. Node interior colors indicate genomic location of genes (yellow: euchromatic regions with normal recombination; gray: euchromatic regions with reduced recombination; blue: heterochomatin). Node border colors denote annotated gene (gray) or NTR (red). Node shape indicates whether a gene is a regulator and/or target (triangles: regulator only; squares: target only; circles: both regulator and target). The node size indicates the number of node connections. Arrows on the edges point to the target. Edges are color coded to show female-specific regulation (red), male-specific regulation (blue) and regulation common to both sexes (black).

There are more NTRs than expected among genes with *cis*-*trans* eQTLs based on the total number of NTRs with eQTLs among the target genes (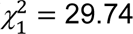, *P* =4.95E-08 in females; 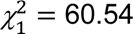, *P* = 7.20E-15 in males) but not the regulatory genes (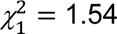, *P* = 0.21 in females; 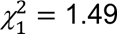, *P* = 0.22 in males). The regulatory genes tend to be located in pericentromeric regions of reduced recombination (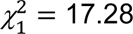, *P* = 3.23E-5 in females; 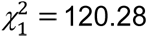, *P* = 0 in males) and target gene locations are enriched for heterochromatin and pericentromeric regions of reduced recombination (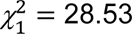, *P* = 9.21E-8 in females; 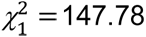, *P* = 0 in males). Regulatory genes with many target genes thus tend to have multiple *cis*-eQTLs in LD near the centromere, and regulate other NTRs both in heterochromatic regions across the genome and euchromatic regions on other chromosomes (Figures 3, S3, S4, S5). The smaller networks with fewer regulators and targets tend to consist of genes in euchromatin in regions of normal recombination (Figures 3, S3, S4, S5; Tables S5C, S5D). Regulatory genes often have many *cis*-eQTLs; a single *cis*-eQTL can regulate multiple target genes; and multiple *cis*-eQTLs within a gene can regulate different target genes. Each gene with at least one *cis*- eQTL may itself be regulated in *trans* by *cis*-eQTLs in one or more upstream genes, and the genes regulated by a focal *cis*-eQTL may themselves have *cis*-eQTLs regulating other genes.

### Genetic Variation in TE Expression

A total of 9% of the *D. melanogaster* genome contains TEs spanning multiple families^32^. Active retrotransposon sequences are present in our RNA-seq libraries. We aligned reads to the RepBase database of known repetitive elements^33^, and quantified TE RNA levels based on normalized read counts. Overall, 1.3% of the RNA-seq reads align to RepBase. The most abundant families of TE sequences were *gypsy*, *copia*, *BEL*, *jockey* and *Mariner*/*Tc1* elements, but all TE families represented in RepBase were detected (Figure 4A, Table S6A).

**Figure 4.**
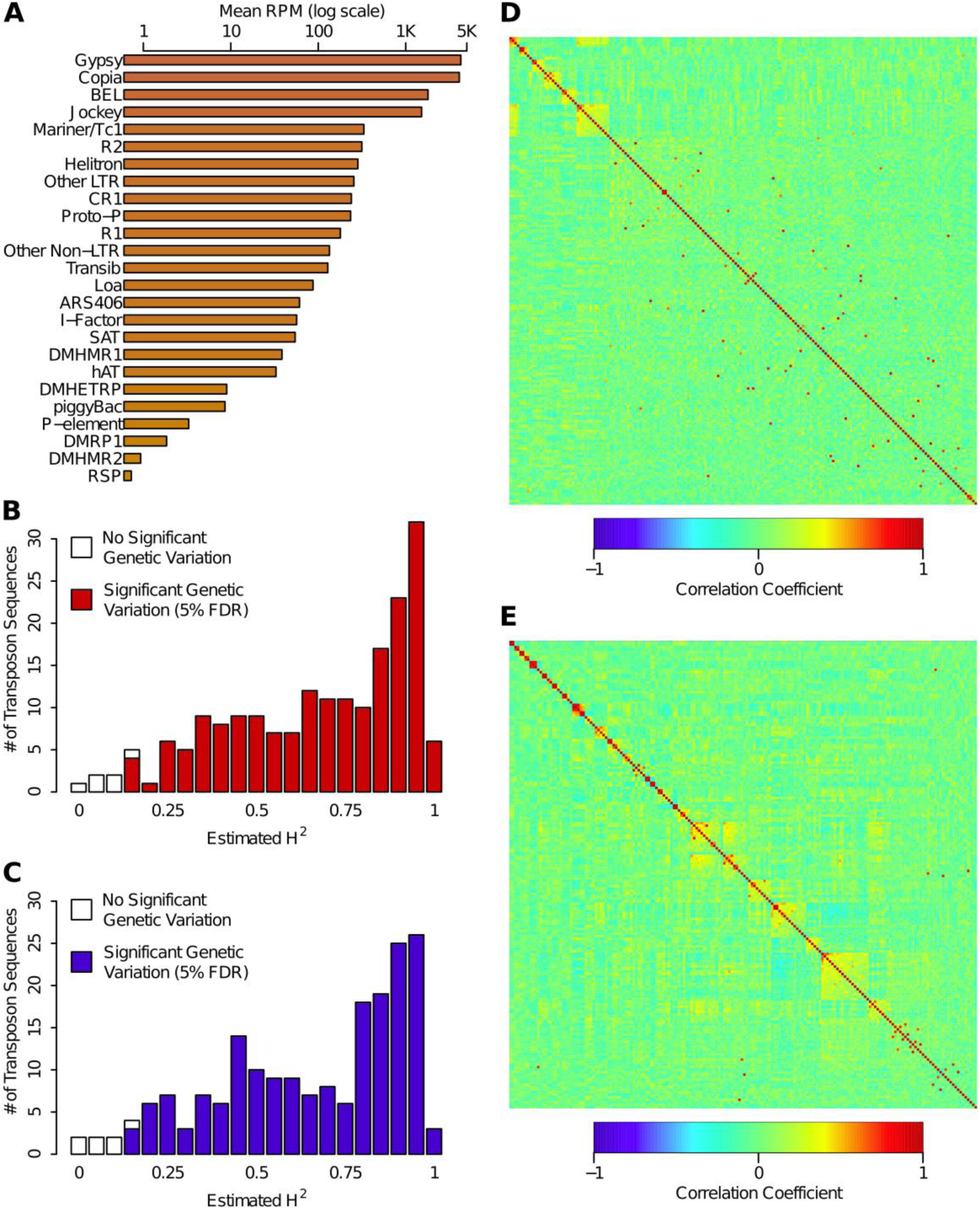
Genetic variation of TE expression in the DGRP. (**A**) Total signal for each TE family, summed over all individual transposon sequences and averaged across all DGRP lines, sex, and replicates. (**B**) Distribution of copy number independent *H*^2^ estimates for TE sequences in females. (**C**) Distribution of copy number independent *H*^2^ estimates for TE sequences in males. (**D**) WGCNA modules of TEs for females. (**E**) WGCNA modules of TEs for males. Heatmaps are depicted as in Figure 1. TE sequences not assigned to any module are included at the bottom right.

Line-specific differences in TE RNA levels can be driven by both differences in underlying copy number^34^ and differences in the rate of transcription per genomic copy. We quantified DNA copy variation for each TE sequence (Table S6B) and used linear models to estimate the percentage of variation in TE expression that arises from differences in copy number (Table S6C). We then partitioned the remaining copy number-independent variation in TE expression between sexes, DGRP lines, the line by sex interaction and residual terms (Table S6C), using FDR ≤ 0.05 as the significance threshold for each term in the analysis. Since the majority (153, 79%) of TEs had a significant sex by line interaction effect, we assessed genetic variation in TE expression for each transposon sequence separately for each sex (Tables S6D, S6E). We observed significant genetic variation in expression for 187 (97%) TE sequences in females (Figure 4B) and 186 (96%) TE sequences in males (Figure 4C). Broad sense heritabilities of TE expression ranged from *H*^2^ = 0.15 – 0.99 in females and *H*^2^ = 0.15 – 0.98 in males (Figures 4B, 4C). Thus, there is host genetic control of expression for most *D. melanogaster* TEs.

We assessed whether different TE sequences had similar patterns of expression across the DGRP lines^28^, separately for males and females (Figures 4D, 4E, Tables S6F, S6G). We found minimal correlation structure in the activity scores of different TEs (Table S6H), with the strongest correlations between pairs of TE sequences from the same family. This suggests that host genetic factors independently affect variation in expression of each TE family.

### TE eQTLs

We mapped eQTLs for each of the TEs with genetically variable expression in females and males (Table S7). We found 54 TEs with significant eQTLs (FDR ≤ 0.05), 36 in females and 39 in males. A total of 20 TE sequences were expressed in both males and females; surprisingly, 16 (18) TE sequences were expressed only in females (males). The number of eQTLs per TE sequence ranged from 1-1,020, with on average more eQTL associations for TEs in males than females (Tables S7A-C). Interestingly, the large numbers of eQTLs associated with some TEs were located in LD blocks in pericentromeric regions and on the *4*^th^ chromosome (Figure S6, Tables S7D, S7E). Many eQTLs for TEs expressed in both males and females overlapped between the sexes, but typically additional eQTLs were present in males. Although there was little clustering of expression patterns of different TE sequences, 202 (1,032) eQTLs were associated with two or more sequences in females (males) (Tables S7F, S7G).

Many eQTLs associated with TE expression were within 1 kb of annotated genes and NTRs. Indeed, 19.8% (17.7%) of TE eQTLs were within 1 kb of NTRs in females (males). Known genes near TE eQTLs were enriched (FDR < 0.05) for GO categories related to regulation of gene expression and protein binding (Table S7H). We next asked to what extent eQTLs associated with gene expression were also associated with expression of TE sequences. We found 1,206 eQTLs associated with 85 genes (37 known genes and 48 NTRs) and 23 TEs in females; and 3,656 eQTLs associated with 166 genes (79 known genes and 87 NTRs) and 30 TEs in males (Figure S7, Table S8). We could thus incorporate variation in TE expression into the *cis*-*trans* gene regulatory network via shared eQTLs (Figure 5). These eQTLs are predominantly located in pericentromeric regions, and the genes they regulate are in pericentromeric regions as well as heterochromatin.

**Figure 5.**
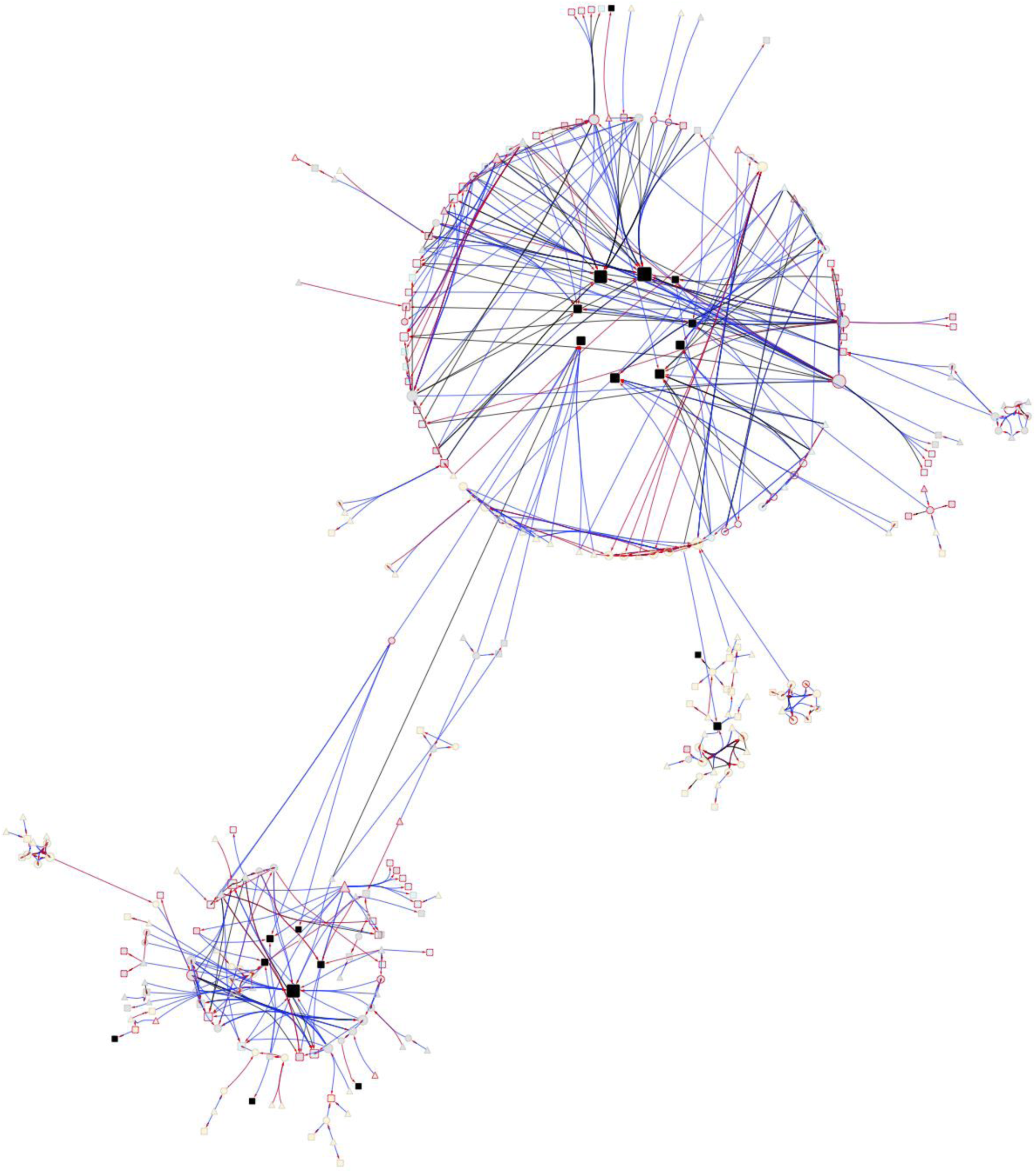
TE genetic regulatory network. Symbols and color-coding are as for Figure 3. Black squares denote TE sequences.

### Genetic Variation in Microbiome Composition

RNA samples extracted from pools of whole flies contain RNA from gut microbial communities, and from microbes on their exoskeleton. We assessed the contribution of microbial sequences to the RNA-seq libraries by aligning reads to a database of candidate microbial genomes (Table S9). *Wolbachia pipientis*, a bacterial endosymbiont that infects ∼50% of the DGRP lines^20^, is the most abundant source of expressed sequence, followed by multiple Acetobacter species and genome assemblies (Figure 6A, Table S9). We estimated the total gene expression from each microbial species in all samples (Table S10A) and partitioned variation in microbial gene expression between sexes, DGRP lines, the sex by line interaction and residual terms, using FDR ≤ 0.05 as the significance threshold (Table S10B). The *H*^2^ of *Wolbachia pipientis* abundance is extremely high (*H*^2^ = 0.972), as expected. We next assessed whether the sum of all non-Wolbachia microbial species is genetically variable after accounting for any Wolbachia effects, and estimated *H*^2^ = 0.595 (Figure 6B, Table S10B). The sex by line interaction for total microbial gene expression was not significant, indicating that total microbial RNA is highly correlated between males and females. We estimated the heritability of gene expression for the 122 non-Wolbachia microbial species, and found that 84 microbial species had significant genetic variation in RNA abundance, with broad sense heritabilities ranging from *H*^2^ = 0.07 – 0.90 (Figure 6C, Table S10B). Microbial species that are likely to colonize the Drosophila gut (Acetobacter and Lactobacillus species) were among those with the highest *H*^2^.

**Figure 6.**
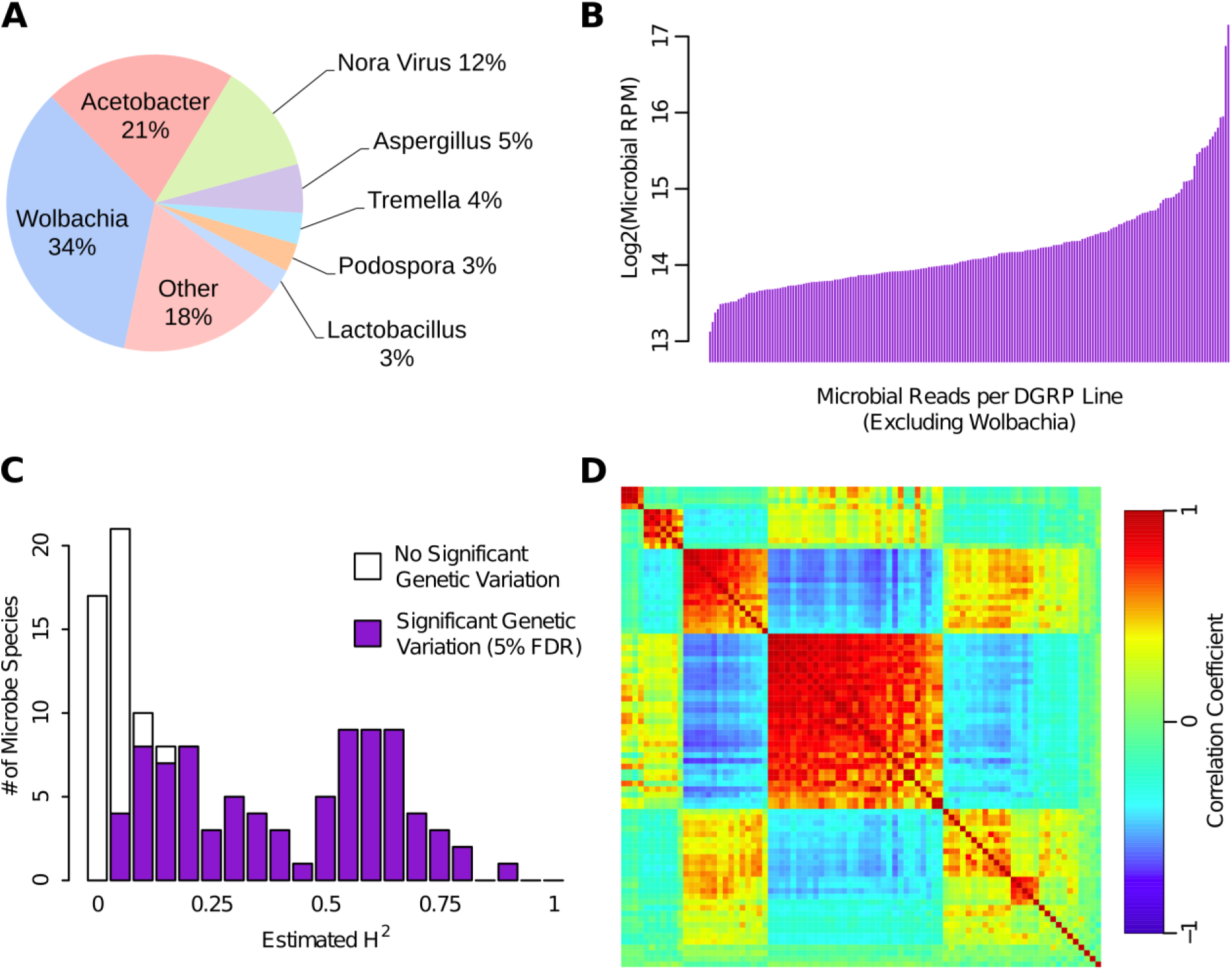
Genetic variation of microbiome composition. (**A**) The proportion of microbiome signal in RNA-seq libraries aligned to species in each genus or viral group. (**B**) Line means of total microbial signal (excluding Wolbachia). (**C**) Distribution of *H*^2^ estimates for individual microbe species. (**D**) WGCNA modules for microbial species. Heatmaps are depicted as in Figure 1. Species not assigned to any module are included at the bottom right.

We used WGCNA^28^ to group species with similar abundance patterns based on the average of male and female line means (Figure 6D, Tables S10C, S10D). We found three groups of strongly correlated species, consisting primarily of the gut-related microbes (Acetobacter and Lactobacillus species), and two additional clusters of microbes primarily consisting of viral and fungal species that are strongly anti-correlated with the abundances of species in the first three clusters. Thus, there is line-specific variation in the microbial communities living in and on DGRP flies. Species which most plausibly colonize the Drosophila gut are largely correlated across lines, with some fluctuation in the relative abundance of Acetobacter versus Lactobacillus species.

### eQTLs for Microbiome Composition

There was little genetic variation in sexual dimorphism for microbial gene expression; therefore, we performed eQTL mapping using the average expression of males and females for each microbial species. Four microbial species and total microbial sequence expression were associated with significant eQTLs (FDR ≤ 0.05) (Table S11A). The sum of all microbial species is associated with one eQTL that maps to an NTR; the expression of *Borrelia coriaceae*, *Acidovorax temperans* and *Podospora anserine* map, respectively, to single eQTLs in *CG2616, CG46301,* and to *cic* and an NTR; and *Leuconostoc pseudomesenteroides* expression maps to 39 variants in or near *GC* and *nSyb* (Table S11A).

We lowered the significance threshold to *P* < 10^−5^ to explore the extent to which common eQTLs may control the expression of multiple microbial species that cluster together based on the WGCNA analysis (Figure 6D). At this threshold, 1,455 eQTLs are associated with 88 microbial species and the sum of all species (Table S11B); 268 variants were associated with expression of more than one microbial species, and five eQTLs were associated with expression of 10 or more microbial species (Table S11C). These data suggest that there is genetic variation in host control of microbial gene expression and that some variants have pleiotropic effects on multiple microbial species.

We assessed whether the genes to which the eQTLs associated with variation in microbial gene expression were enriched for GO categories (FDR ≤ 0.05). The most highly enriched Biological Process GO terms were related to development and morphogenesis, including development and function of the nervous system (Table S11D).

### Gene Expression and Complex Traits

To examine the relationship between variation in gene expression and variation in organismal quantitative trait phenotypes, we chose 11 quantitative traits with published phenotypic data (chill coma recovery time and startle response^19^; starvation resistance^20^; day and night sleep bout number, day and night total sleep duration, and total waking activity^35^; food consumption^36^; male aggression^37^; phototaxis^38^); and additionally measured five metabolic traits (levels of free glucose, glycogen, free glycerol, triglyceride and protein) and three metrics of body size (body weight, thorax length, thorax width). All traits were quantified in the same laboratory under the same culture conditions used in this study. The line means for all traits are given in Table S12; quantitative genetic analyses of the metabolic and body size traits are given in Table S13; and the most significant associations (*P* < 10^−5^) from GWA analyses (separately for males and females) for these quantitative traits based on the 200 lines for which we have gene expression data are in Table S14.

We first assessed whether variants associated with all organismal traits were enriched for eQTLs, as found in human studies^6,7,9,14,15^. We found no enrichment of *cis*-eQTLs (*P* = 0.13 in females and *P* = 0.71 in males), *trans*-eQTLs (*P* = 0.98 in females and *P* = 0.28 in males) or all eQTLs (*P* = 0.94 in females and *P* = 0.23 in males) among top GWA hits in either sex. Many top GWA hits as well as eQTLs map to regions greater than 1kb from any gene, and may indicate novel regulatory regions.

We next performed transcriptome wide association studies (TWAS) for individual genetically variable transcripts for gene expression, TE sequences and microbial species, for each of the 18 (19) genetically variable organismal phenotypes in females (males). We found several significant (Benjamini-Hochberg FDR < 0.05) associations of transcripts with organismal phenotypes (Table S15). These associations include a known noncoding RNA (*CR46032*) with male aggression, two NTRs with male waking activity, *Gbs-70E* with free glucose in both sexes, *AkhR* with starvation resistance in males and females, and *Acidovorax temperans* with male aggression (Table S15).

## Discussion

Deep RNA sequencing gives accurate estimates of gene expression of annotated genes and can implicate novel non-coding RNAs and their regulatory interactions with annotated genes. lncRNAs are operationally defined as encoding transcripts > 200 bp with no significant protein-coding potential^21–24,39,40^. We have identified 4,282 novel transcribed regions, most of which are likely lncRNAs, increasing the total number of *D. melanogaster* lncRNAs nearly threefold: from 2,366^40^ to 6,648. These lncRNAs are unlikely to be artifacts since the majority are genetically variable, and they are not randomly distributed in the genome but are preferentially located in heterochromatic regions and in pericentromeric euchromatin bordering heterochromatin. Thus, there is genetic variation in heterochromatic gene expression, thought to be largely transcriptionally silent^41^. These heterochromatic and pericentromeric lncRNAs are regulated by pericentromeric *cis*-eQTLs as well as a large number of *trans*-eQTLs dispersed throughout the euchromatic genome. Genes associated with eQTLs with both *cis*- and *trans*- effects form sex-specific networks of regulator and target genes, the largest of which is enriched for lncRNA target genes in heterochromatin and regulator and target genes in pericentromeric euchromatin. The considerable overlap between eQTLs associated with lncRNAs in the large networks and TE expression recruits TEs to the network. We do not know where the TE sequences with genetically variable expression are integrated in the genome; however, heterochromatin is composed of largely silenced TE repeats^41^, raising the possibility that TEs in heterochromatin are subject to the same regulation as other heterochromatic genes. Further work is needed to confirm the regulatory networks derived from naturally occurring genetic variation and determine the regulatory mechanism(s) through which the lncRNAs act^22,24,39,40,42,43^.

The first step in systems genetic analysis is to identify eQTLs associated with both gene expression and organismal quantitative traits, for which variation in gene expression is correlated with variation in the organismal phenotypes^2,5,8^. We did not find any such trios, although we did find interesting transcript-trait associations. This may be because our sample size is adequate to detect eQTLs but not QTLs affecting organismal traits, which have smaller effects; because eQTLs need to be mapped in tissues relevant to the organismal trait; and because there are non-linear (epistatic) relationships between QTLs for both transcripts and organismal phenotypes. The complex and highly connected *cis-trans* regulatory networks suggest that higher order interactions need to be accommodated in systems genetic modeling, at least at the level of gene expression.

## Methods

### Drosophila lines

We used 200 inbred, sequenced DGRP lines^19,20^, established by 20 generations of full sib inbreeding from gravid females collected at the Raleigh, NC USA Farmer‟s Market. Genome sequences of the lines were obtained previously using the Illumina platform with an average of coverage of 27×. A total of 4,565,215 molecular variants (3,976,011 single/multiple nucleotide polymorphisms (SNPs/MNPs), 169,053 polymorphic insertions (relative to the reference genome), 293,363 polymorphic deletions and 125,788 polymorphic microsatellites) segregate in the DGRP.

### Sample collection

All lines were reared on cornmeal-molasses-agar medium at 25°C, 60–75% relative humidity and a 12-hr light-dark cycle at equal larval densities. We collected two replicates of 25 females and 30 males per line, for a total of 800 samples. We used a strict randomized experimental design for sample collection. We collected mated 3-5 day old flies between 1-3 pm. We transferred the flies into empty culture vials and froze them over ice supplemented with liquid nitrogen, and sexed the frozen flies. The samples were transferred to 2.0 ml nuclease-free microcentrifuge tubes (Ambion) and stored at −80°C until ready to process.

### RNA sequencing

Total RNA was extracted with QIAzol lysis reagent (Qiagen) and the Quick-RNA MiniPrep Zymo Research Kit (Zymo Research). Ribosomal RNA (rRNA) was depleted from 5 ug of total RNA using the Ribo-Zero^TM^ Gold Kit (Illumina, Inc). Depleted mRNA was fragmented and converted to first-strand cDNA using Superscript III reverse transcriptase (Invitrogen). During the synthesis of second strand cDNA, dUTP instead of dTTP was incorporated to label the second strand cDNA. cDNA from each RNA sample was used to produce barcoded cDNA libraries using NEXTflex™ DNA Barcodes (Bioo Scientific, Inc.) with an Illumina TruSeq compatible protocol. Libraries were size-selected for 250 bp (insert size ∼130 bp) using Agencourt Ampure XP Beads (Beckman Coulter, Inc.). Second strand DNA was digested with Uracil-DNA Glycosylase before amplification to produce directional cDNA libraries. Libraries were quantified using Qubit dsDNA HS Kits (Life Technologies, Inc.) and Bioanalyzer (Agilent Technologies, Inc.) to calculate molarity. Libraries were then diluted to equal molarity and re-quantified. A total of 50 pools of 16 libraries were made, again randomly assigning samples to each pool. Pooled library samples were quantified again to calculate final molarity and then denatured and diluted to 14pM. Pooled library samples were clustered on an Illumina cBot; each pool was sequenced on one lane of Illumina Hiseq2500 using 125 bp single-read v4 chemistry.

### RNA sequence analysis

Barcoded sequence reads were demultiplexed using the Illumina pipeline v1.9. Adapter sequences were trimmed using cutadapt v1.6^44^ and trimmed sequences shorter than 50bp were discarded from further analysis. Trimmed sequences were then aligned to multiple target sequence databases in the following order, using BWA v0.7.10 (MEM algorithm with parameters ‘-v 2 –t 4’)^45^: (1) all trimmed sequences were aligned against a database containing the complete 5S, 18S-5p8S-2S-28S, mt:lrRNA, and mt:srRNA sequences to filter out residual rRNA that escaped depletion during library preparation; (2) remaining sequences were then aligned against a custom database of potential micriobiome component species (see below) using BWA; (3) sequences that did not align to either the rRNA or microbiome databases were aligned to all *D. melanogaster* sequences in RepBase^33^. The remaining sequences that did not align to any of the databases above were then aligned to the *D. melanogaster* genome (BDGP5) and known transcriptome (FlyBase v5.57) using STAR v2.4.0e^46^. Libraries with fewer than 5 million reads uniquely aligned to the *D. melanogaster* reference genome were re-sequenced to achieve sufficient read depth.

### Generation of microbiome database

We first performed a preliminary alignment of RNA-seq reads by filtering only rRNA sequences, and then aligning directly to the *D. melanogaster* genome using the tools and parameters described above. Sequences that did not align to the rRNA database or *D. melanogaster* reference genome were then analyzed with Trinity v2.1.1 to perform *de novo* assembly of longer sequences from the short reads. Assembled sequences > 1kb in length were then searched against the refseq_genomic database (downloaded from NCBI on 1/27/16) using BLAST. We then compiled a list of all refseq genomes that were found as a top BLAST hit for at least two assembled sequences. We compiled all fasta files for each of these refseq genomes into a single database for alignment with BWA.

### Genotype validation

To validate the DGRP line assigned to each RNA-seq sample, we identified single nucleotide polymorphisms (SNPs) from the RNA-seq reads that aligned to the *D. melanogaster* reference genome using STAR as described above. We retained only those SNP calls covered by at least 3 reads and at least 75% of all reads supporting the major genotype (note that DGRP lines are inbred and therefore the majority of SNPs are homozygous). This filtering process produced >400k usable SNPs per sample, primarily located in transcribed regions of the genome. We then performed two validation tests of the DGRP line assigned to each sample *X* by comparing to the previously published genotype calls for each DGRP line (http://dgrp2.gnets.ncsu.edu/data/website/dgrp2.tgeno20). First, we computed the “line mean error” (LME) for each line as follows: given the set of homozygous SNPs from line *Y* that have sufficient coverage (described above) in sample *X*, LME(*X*,*Y*) = # of mismatching SNPs / total # of comparable SNPs. We confirmed that for each sample *X*, the DGRP line *Y_lab_* labeled for that samples produced the minimum value of LME(*X*,*Y*) as compared to all other possible line assignments *Y_alt_*, and further confirmed that LME(*X*,_Ylab_) was below 1%. Second, we performed competitive tests between the labeled line *Y_lab_* and each possible alternate line *Y_alt_.* Under this test, we considered only the SNPs that are homozygous for different genotypes in *Y_lab_* and *Y_alt_* (*i.e.*, only the segregating SNPs for the two lines) and which have sufficient coverage in sample *X*. We then computed the “line error ratio” (LER) = # of SNPs matching *Y_lab_* /# of SNPs matching *Y_alt_*. We confirmed that for each sample *X*, the lowest LER for any *Y_alt_* was > 1 (*i.e.*, the majority of SNP calls always supported the labeled line compared to any alternative line).

### Inference of novel transcripts

We constructed a *de novo* transcriptome for each individual sample by inputting the RNA-seq reads aligned to the *D. melanogaster* reference genome into Cufflinks v2.2.1^47^. We also considered the novel transcribed regions (NTRs) identified in a previous study based on unstranded pooled RNA sequencing of the DGRP lines^27^. However, the previously published data do not provide strand-specific signal, while our current RNA-seq data uses a strand-specific library preparation. Therefore, we reassigned the strand for each of the previously published NTRs that was supported by the greater number of total aligned reads across all samples. We then merged all *de novo* sample transcriptomes and the previously published NTRs using the cuffmerge tool included with Cufflinks v2.2.1, then removed all merged transcript models with any exon overlapping on the same strand any exon in the known *D. melanogaster* transcriptome. We defined the known transcriptome here as all gene models in FlyBase v5.57 plus all subsequently added gene models in FlyBase v6.11 to account for recently discovered lncRNA sequences. Thus, the final output of this analysis was a set of NTRs constructed from both our current RNA-seq data and previously published pooled RNA-seq data that do not overlap any known gene exons on the same strand.

### Gene expression estimation

Read counts for individual microbial species were computed as all reads aligning to any sequence in any genome for any strain of that species. Reads aligning to multiple species were ignored for individual species read counts. We also aligned microbiome-aligning reads to the *D. melanogaster* genome, and removed all reads that aligned to both microbial and *D. melanogaster* sequences before computing read counts, to account for several domains which are highly conserved between microbial and metazoan species. Read counts were computed for transposon sequences by computing the number of reads uniquely aligned to each transposon sequence in RepBase. Highly homologous sequences were grouped together for computing transposon read counts. Read counts were computed for known and novel gene models using HTSeq-count^48^ with the „intersection-nonempty‟ assignment method. Tabulated read counts for each expression feature type (microbiome, transposon, endogenous genes) were then normalized across all samples using EdgeR^49^ as follows. First, genes with low expression overall (<10 aligned reads in >75% of the libraries) were excluded from the analysis. Library sizes were re-computed as the sum of reads assigned to the remaining genes, and further normalized using the Trimmed Mean of M-values (TMM) method^50^. At this point, we retained only genes (known or novel) whose expression in both biological replicates was above an empirical threshold in more than 200 line-sex combinations (400 samples total). This criterion retains genes expressed in only one sex. The threshold was determined by fitting all log2 transformed FPKM expression data points using a 2-component Gaussian mixture model and finding the expression value (FPKM = 0.280263) where the posterior probability of being in the lower expression component is 0.95. Genes on chrU and chrUextra were also removed. We further adjusted transposon expression estimates to account for differences in transposon copy number across lines by fitting a linear model: RNA ∼ DNA + *ε*, where RNA = the normalized log2(RNA-seq read count); and DNA = normalized log2(DNA read count) derived from the previously published DNA-seq data for each DGRP line^20^. After fitting the linear model for each transposon sequence, *ε* estimates the relative transcription rate in each line independent of copy number, and was used as the adjusted transposon expression for all subsequent analysis. We further adjusted endogenous gene expression values by estimating and removing the effect of alignment bias resulting from higher rates of non-reference variants clustering in some lines. We computed the alignment bias score *A*(*g*,*L*) defined as the number of non-reference nucleotides per kb in all exons of gene *g* in DGRP line *L*, based on the previous map of genomic variation in the DGRP^20^. We then fit a linear model for each endogenous gene: *Y* = *A* + *ε*, where *Y* is the normalized expression profile for gene *g* after the read counting and EdgeR normalization described above. After fitting these linear models, *ε* represents the alignment bias-corrected expression, and was used as the normalized gene expression in all subsequent analysis.

### Genetics of gene expression

For each class of expression features (endogenous genes, transposons, microbiome), we fit mixed-effect models to the gene expression data corresponding to: *Y* = *S* + *W* + *W*×*S* + *L* + *L*×*S* + *ε*, where *Y* is the observed log2(normalized read count), *S* is sex, *W* is Wobachia infection status, *W*×*S* is Wolbachia by sex interaction, *L* is DGRP line, *L*×*S* is the line by sex interaction and *ε* is the residual error. We also performed reduced analyses (*Y* = *W* + *L* + *ε*) independently for males and females. We identified genetically variable transcripts as those that passed a 5% FDR threshold (based on Benjamini-Hochberg^51^ corrected *P*-values) for the *L* and/or *L*×*S* terms. We computed the broad sense heritabilities (*H*^2^) for each gene expression trait separately for males and females as 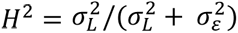, where 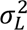 and 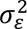 are, respectively, the among line and within line variance components.

### Clustering by genetic correlation

For each feature type (microbiome, transposons, endogenous genes), we clustered line means using the WGCNA R package v1.51^28^ as follows. Only genes with sufficient average expression (Log2 FPKM > 0) and genetic variance (line mean variance > 0.05) were considered in these analyses. First, the Pearson correlation coefficient for every pair of line means, the soft-power threshold was computed using the pickSoftThreshold function, and used to convert the correlation matrix to an adjacency matrix with approximately scale-free connectivity. The adjacency matrix was then converted to a dissimilarity matrix based on the topological overlap map^28^. Expression features were then clustered using hierarchical clustering (hclust function) based on the dissimilarity matrix, and split into distinct modules using the cutreeDynamic with deepSplit=4 and minClusterSize=20 (for endogenous gene expression, minClusterSize=4 was used for microbiome and transposon clustering). Module eigengenes were computed for each cluster, and highly similar clusters were combined using the mergeCloseModules function with cutHeight = 0.25. Expression features assigned to module 0 (insufficient similarity) were discarded. Modules consisting of >1,000 features were also discarded as insufficiently split into distinct modules. For each expression feature, the degree was computed as the overage topological overlap with all other features assigned to the same module. The average degree of each module was computed as the mean degree across all features in the module. Modules were sorted by average degree, such that module 1 has the highest average degree in each analysis.

### Gene set enrichment analyses

Lists of known gene IDs (FlyBase FBgn accessions) were uploaded to FlyMine^29^ or Panther^52^ for functional enrichment. For analysis of gene lists from WGCNA clusters, the list of known genes input to WGCNA was used as the background set, to correct for any biases inherent to highly heritable expression patterns in general.

### Expression QTL (eQTL) mapping

For each gene expression feature, we performed eQTL analysis as previously described^26^. Briefly, we adjusted mean expression values in each sex for fixed effects of Wolbachia infection status, five major polymorphic inversions (*In2L(t)*, *In2R(NS)*, *In3R(P)*, *In3R(K)*, *In3R(Mo)*), and the first 10 principal components of the genetic relatedness matrix of all DGRP lines using a linear model. We mapped QTLs for the adjusted line means using PLINK^53^ against 1,932,427 SNPs with major allele frequency > 0.05 and missing genotypes in fewer than 25% of the 200 DGRP lines profiled by RNA-seq. We computed FDR of eQTL calls by comparing observed eQTL *P*-value distributions to those obtained from running PLINK on 100 permutations of the observed line means for each expression feature. At any given *P*-value cut-off *X*, the estimated false positive rate of eQTLs for a specific gene expression feature is the average number of eQTLs with *P*-value < *X* across all permutations. The FDR at the same *P*-value is then computed as the estimated false positive rate divided by the number of eQTLs with *P*-value < *X* in the observed data. Using this formulation of FDR, we identified the unadjusted *P*-value cut-off corresponding to 5% FDR for each gene expression feature. No further model selection was performed, however we classified eQTLs as being either *cis*-eQTLs (within 1kb of the gene body for the associated expression feature) or *trans*- eQTLs (> 1 kb of the gene body).

### Construction of eQTL networks

We then constructed regulatory eQTL networks based on individual SNPs which were called as both *cis*- and *trans*-eQTLs for multiple expression features. Specifically, we assign a directed edge *X* → *Y* if there is at least one variant that is both a *cis*-eQTL for gene *X* (defined as within 1 kb of gene *X*) and a *trans*-eQTL for gene *Y* at 5% FDR. We then broke all loops in the regulatory network for each sex by dropping the edge in each loop with the highest minimum *P*-value from all associated SNPs to create a directed, acyclic network.

### Quantitative traits

We retrieved phenotypic data documented from previous publications on the same fly lines for male aggression^37^; chill coma recovery time and startle response^19^; food consumption^36^; phototaxis^38^; sleep traits^35^ (day and night bout number, day and night total sleep duration, total waking activity); and starvation resistance^20^.

To measure body weight and size, we collected 10 replicates of 10 flies per line and sex into pre-weighed 1.7 ml tubes, and weighed and flash froze them for downstream analyses. Virgin flies were used to avoid body weight variation due to variation in egg production. In addition we measured thorax length and thorax width as metrics for body size.

Frozen flies were homogenized in 250 μL Dulbecco‟s phosphate-buffered saline, and after gentle centrifugation supernatants were collected for measurements of free glucose, glycogen, free glycerol, triglyceride and total protein (further diluted 10 fold). For free glucose and glycogen, samples were denatured at 95°C for 25 minutes to prevent glycogenolysis. Measurements were performed following protocols provided by the Glycogen Colorimetric/Fluorometric Assay Kit (BioVision Inc.). For free glycerol and triglyceride, we used the Serum Triglyceride Determination Kit (Sigma Aldrich Inc.), and incubated samples with the Triglyceride Reagent for 1 hour at 37°C. For total protein measurement, we used the Qubit Protein Assay Kit (Thermo Fisher Scientific Inc.).

### Quantitative trait genetic parameters

We used mixed model, factorial ANOVAs (*Y* = *S* + *L* + *L*×*S* + *Rep*(*L*) + *S*×*Rep*(*L*) + *ε*, to partition variation of the quantitative traits between sexes (*S*), DGRP lines (*L*) and replicate vials within lines (*Rep*). Broad sense heritabilities were estimated as 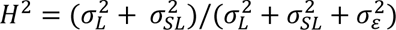, where 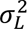, 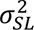 and 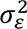 are, respectively, the among line, sex by line and within line variance components.

### eQTL-GWA enrichment analysis

We performed GWA analyses for all quantitative traits, separately for females and males. All phenotypes (line means) were first adjusted for the effect of Wolbachia infection and major polymorphic inversions using a linear model. The residuals (plus the intercept) from this analysis were then used as phenotype in a linear mixed model to test for the effect of each common variant individually, while adjusting for sample structure using a genomic relationship matrix (GRM), as implemented in GCTA-MLMA^55^. The GRM was built as 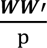 where ***W*** is a matrix of centered and scaled genotypes for the 200 lines and p is the total number of genetic variants.

For each trait and sex, variants with *P* < 10^−5^ were retained for downstream analysis. We then combined the lists of variants associated with each trait, separately for females and males, to obtain a single list of unique variants (*i.e.*, no duplicates) associated with any of the traits of interest. The enrichment analysis proceeded as described in Ref. 14, within each sex. Briefly, GWAS hits were divided into minor allele frequency bins of width equal to 0.05. Then, an equal number of common variants (which may or may not have included actual GWAS hits) per bin were sampled at random and the overlap with eQTLs was calculated. This procedure was repeated 10,000 times and an empirical *P*-value for the enrichment was calculated as the number of replicates where the overlap between randomly sampled variants and eQTLs was greater than or equal to the observed overlap between GWAS hits and eQTLs over the total number of replicates.

### Association of expression and quantitative traits

A transcriptome-wide association study (TWAS), *i.e.,* regressing the phenotype on each gene‟s expression level, was performed for each sex separately for each quantitative trait. We developed a method that accounts for structure present in the transcriptome due correlations between transcripts. This was achieved by fitting a linear mixed model of the type: **y = 1*μ + wβ +* t + e**, where **y** = n-vector of mean phenotypic values for n lines, *μ* = fixed population mean effect, **w** = n-vector of the tested gene‟s centered and scaled expression level, *β* = fixed effect of the gene, **t** = n-vector of random transcriptomic line effect (**t** ∼*N*(0, ***T****σ^2^_t_*)), and **e** = n-vector of random error (**e** ∼*N*(0, ***I****σ^2^_e_*)).

The key term in the model that accounts for sample structure is **T**, the transcriptomic relationship matrix (TRM). The TRM was computed as 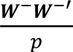, where ***W^−^*** is a matrix of centered and scaled gene expression levels for the 200 lines, excluding the gene tested to maximize the power to find an association^54^, and *p* is the total number of genes. The TRM in TWAS has similar role to the GRM in GWAS.

The effect of each gene’s expression level on the phenotype was tested using a Wald test of the form 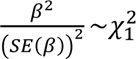. Raw *P*-values and Benjamini-Hochberg FDR-corrected *P*-values^51^ were computed.

The phenotypes were adjusted for the effects of Wolbachia and major polymorphic inversions as described in the previous section. Because the phenotypes were adjusted, we did not adjust gene expression in this analysis to avoid spurious associations due to adjustment on both sides of the equation.

We also performed similar associations of quantitative traits with TEs and microbial gene expression, using the same models as for TWAS but substituting TE and microbial expression for gene expression levels. Quantitative trait phenotypes were adjusted for the effects of Wolbachia and major polymorphic inversions but the TE and microbial expression data were not. The TE analysis was performed for males and females separately, while sex-pooled microbe expression data was used with female or male quantitative trait phenotypes since microbial gene expression was not sex-specific.

## Data Availability

All RNA sequence data have been deposited in GEO (accession GSE117850). The DGRP lines are available from the Bloomington Drosophila Stock Center.

## Acknowledgements

This work was supported by National Institutes of Health grants R01 AA016560, R01 AG043490 and U01 DA041613 to T. F. C. M and R. R. H. A. and Genomic Selection in Animals and Plants (GenSAP) funded by The Danish Council for Strategic Research to T. F. C. M. and F. M.

## Author Contributions

L. E., S. Z., W. H., F. M., R. R. H. A. and T. F. C. M. wrote the manuscript. L. E., S. Z., W. H., F. M. and T. F. C. M. analyzed the data. M. A. C., R. L., G. A., M. S. G., J. M., G. S. A. and L. T performed the research.

## Author Information

The authors declare that no competing interests exist.

---

**Figure S1.**
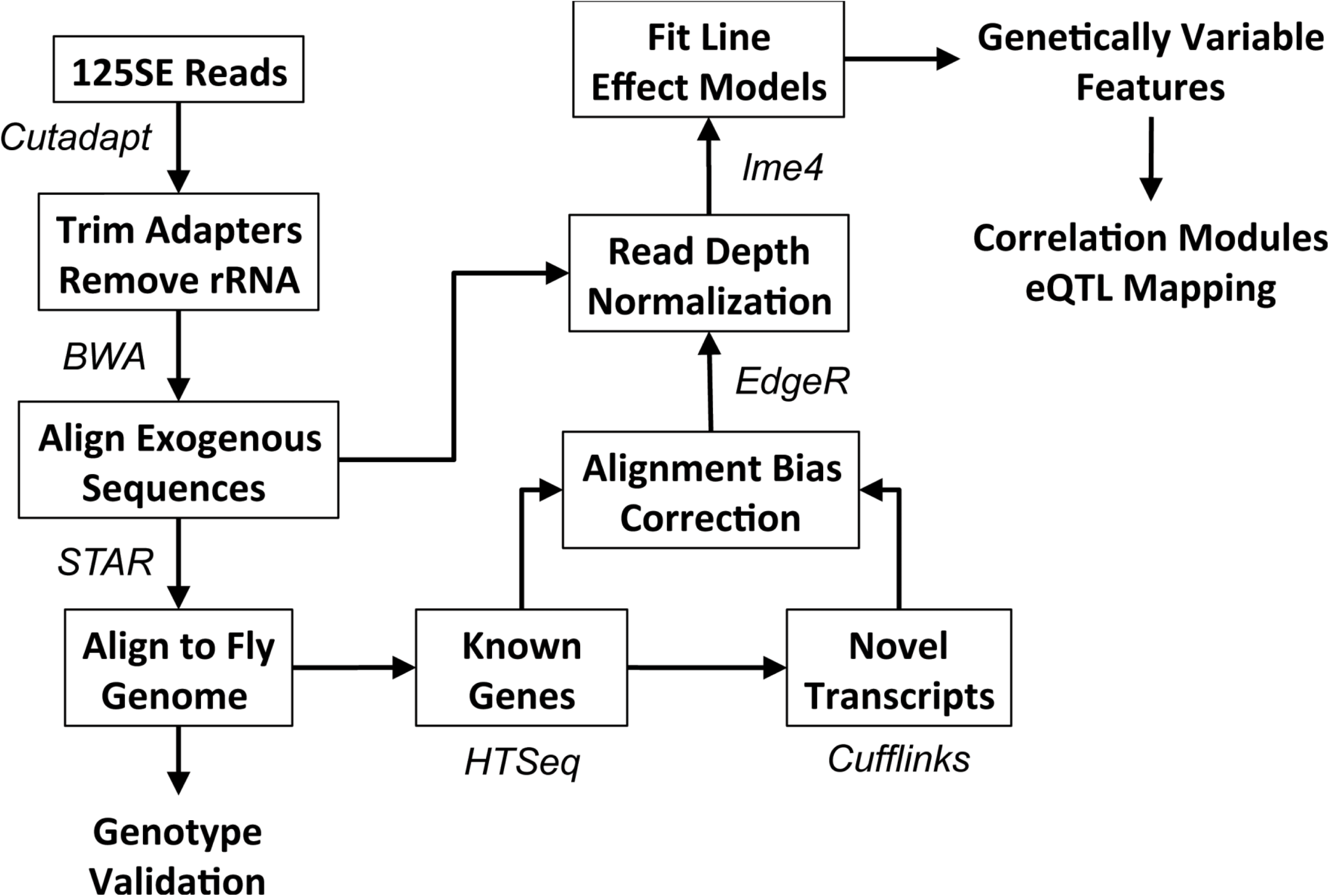
Schematic of the bioinformatics pipeline used for RNAseq analysis.

**Figure S2.**
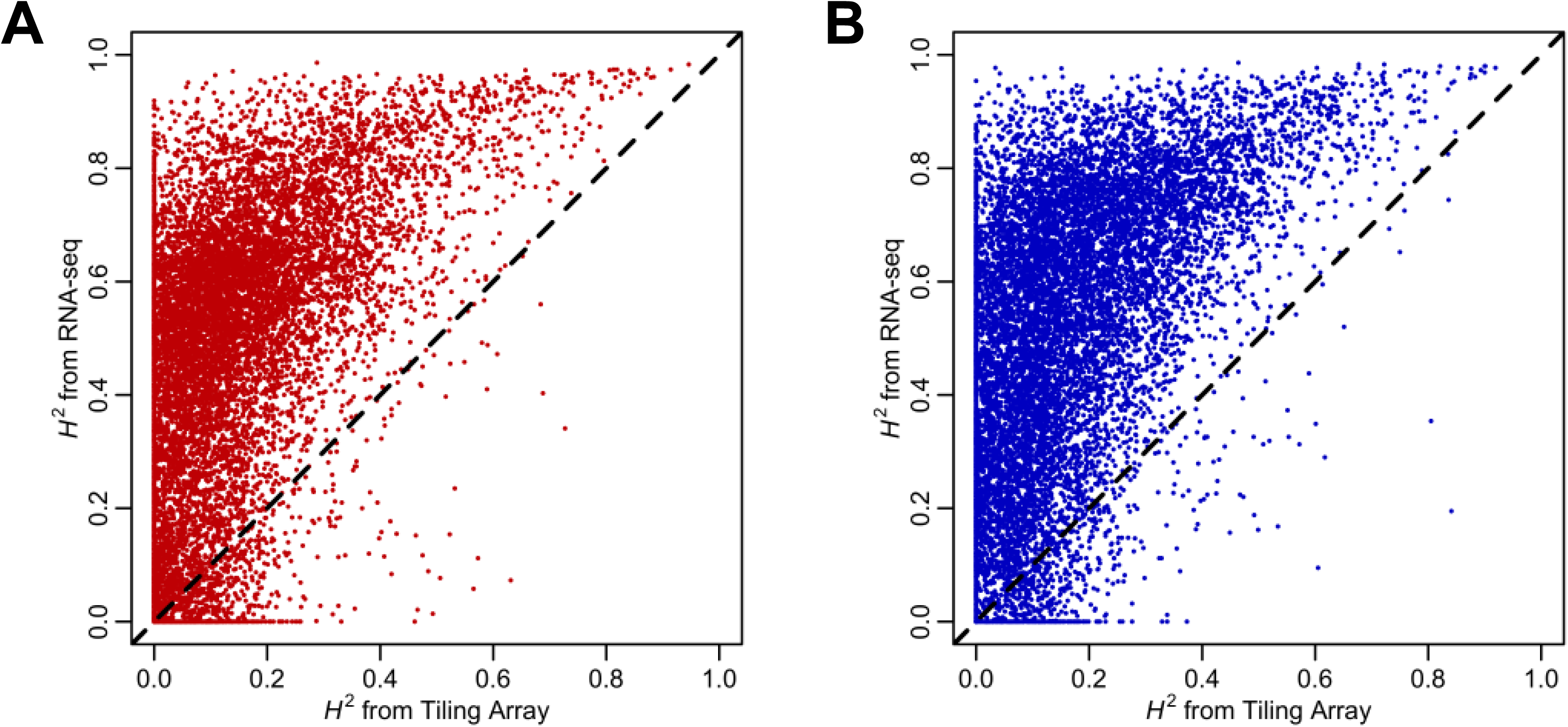
Comparison of RNA-seq and tiling arrays. Scatter plots show the broad-sense heritability (*H*^2^) estimates from RNA-seq in this study compared to tiling array data^26^. (A) Female gene expression (*r* = 0.56, *P* < 1E-15). (B) Male gene expression (*r* = 0.55, *P* = 1E-15).

**Figure S3.**
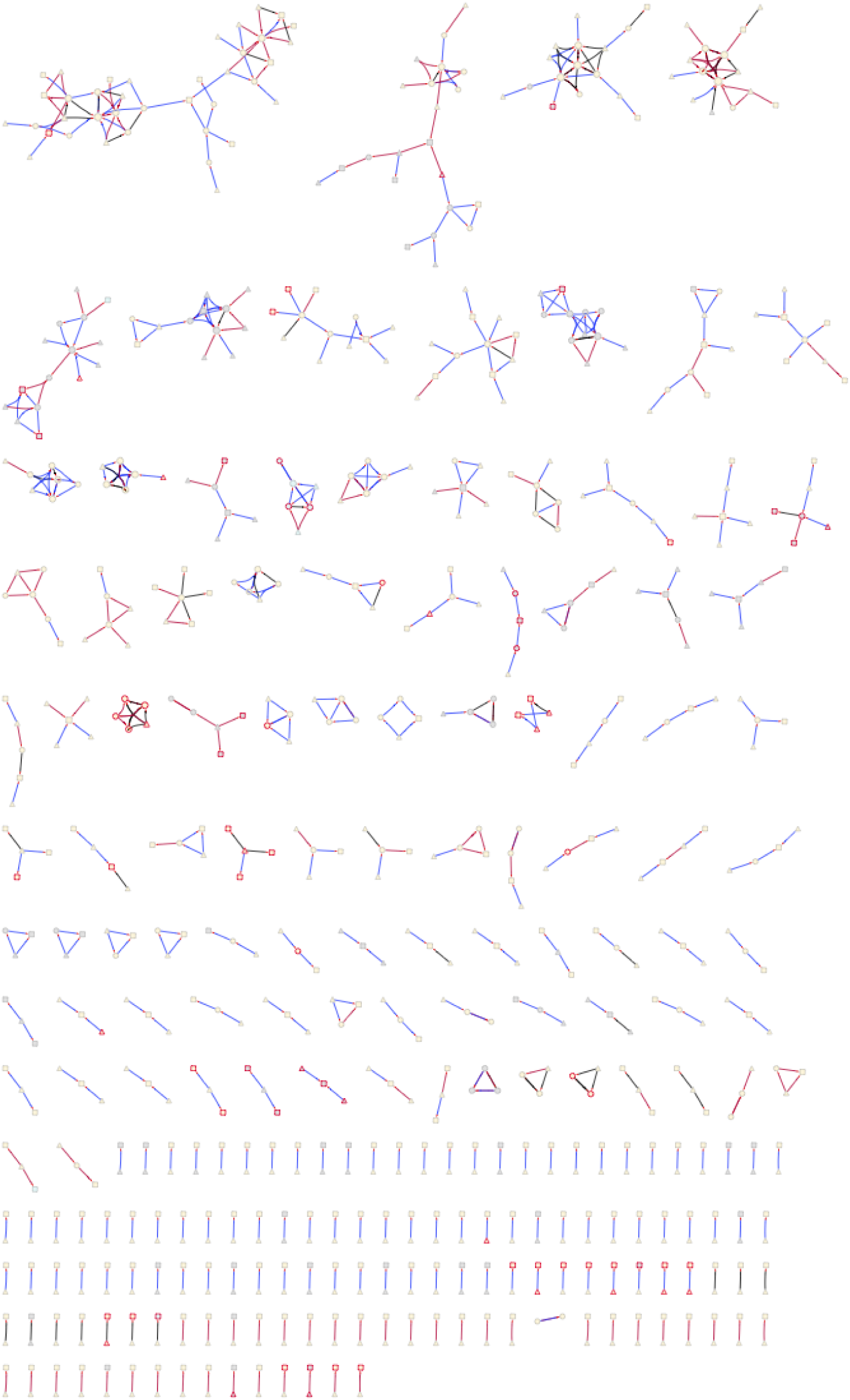
Small *cis*-*trans* eQTL genetic networks in females and males. Symbols and color coding are as in Figure 3.

**Figure S4.**
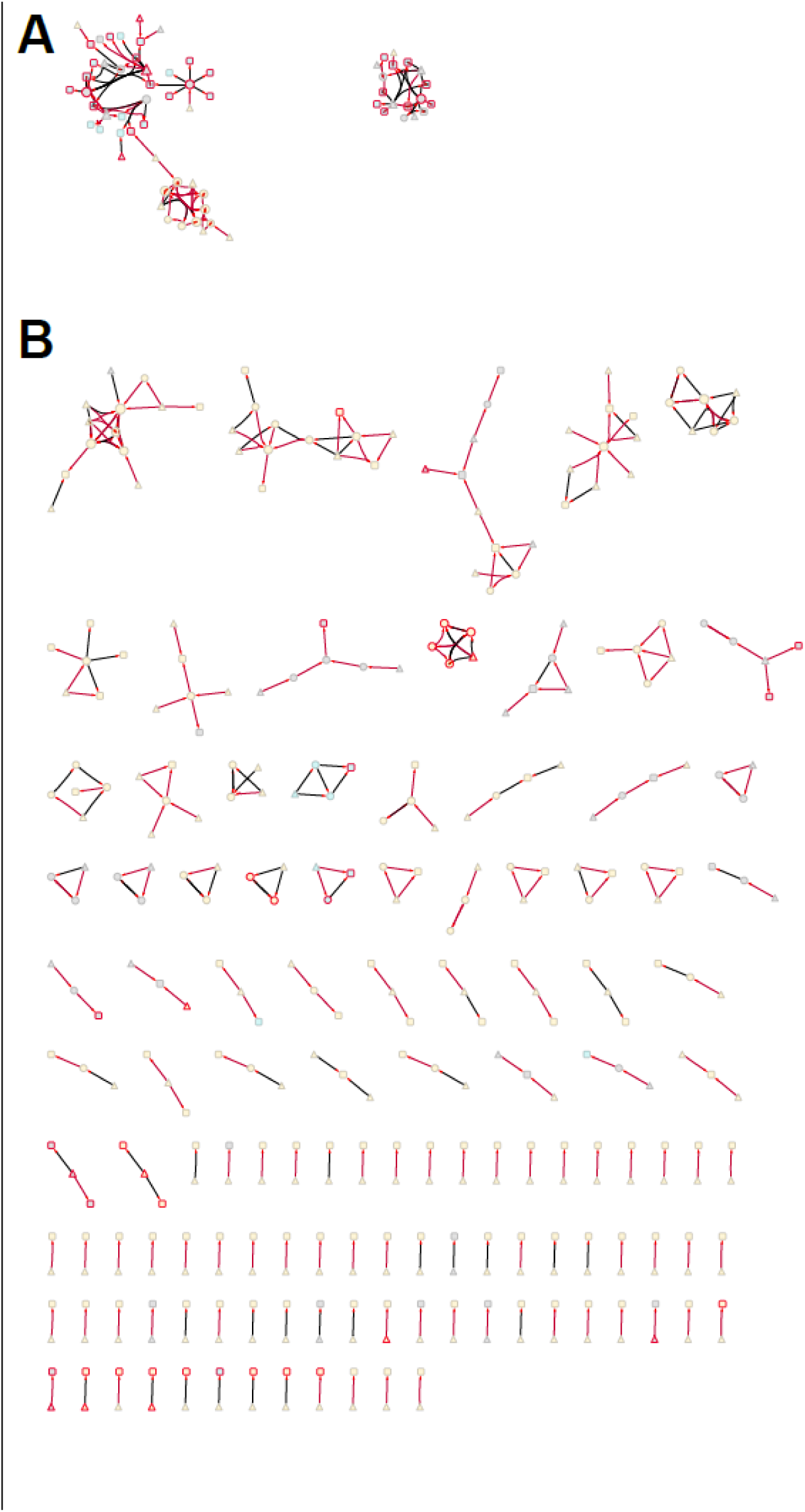
Female *cis*-*trans* eQTL genetic network. Symbols and color-coding are as for Figure 3. (**A**) Network 1 and 2. (**B**) Other networks.

**Figure S5.**
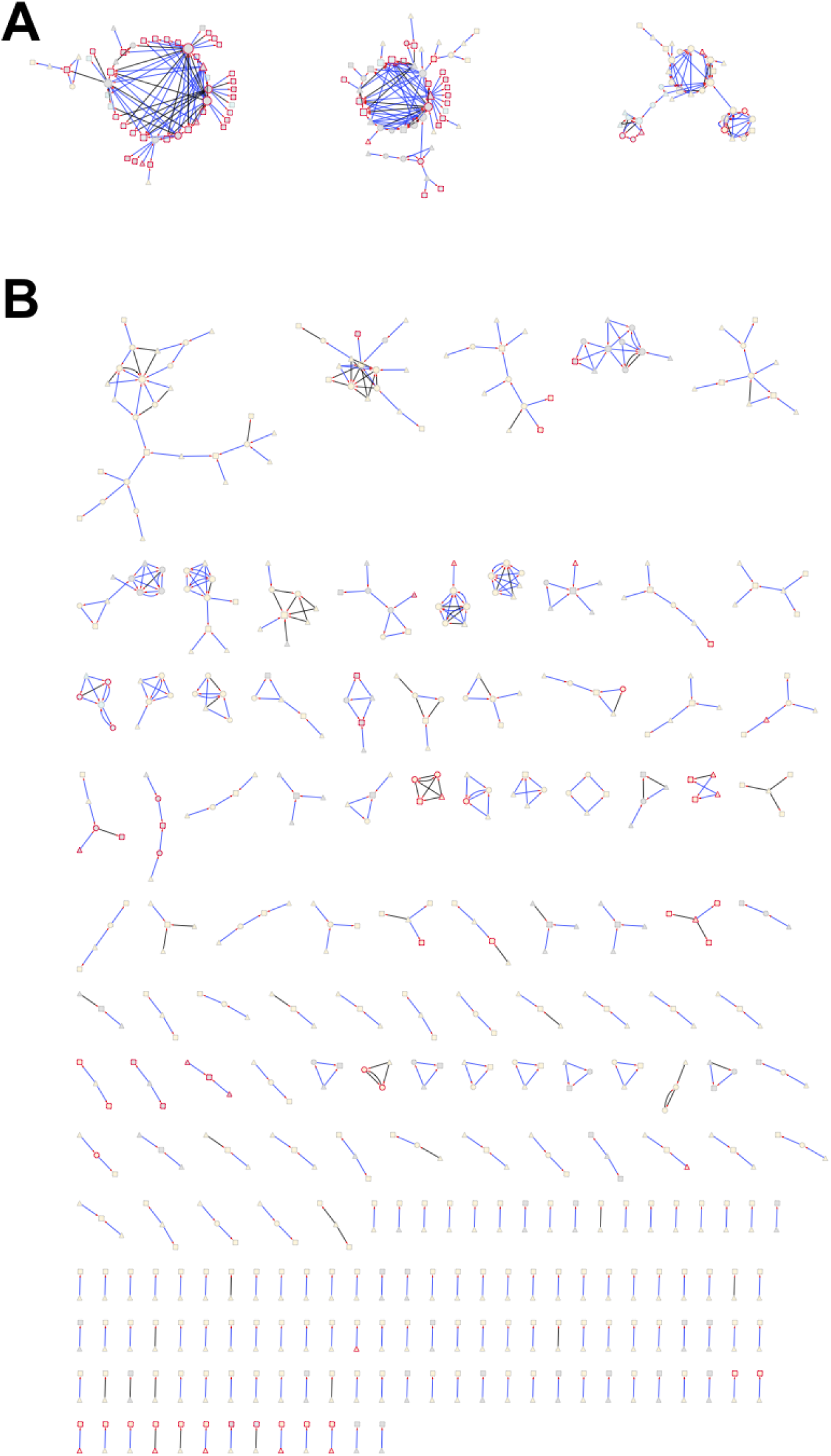
Male *cis*-*trans* eQTL genetic network. Symbols and color-coding are as for Figure 3. (**A**) Networks 1-3. (**B**) Other networks.

**Figure S6.**
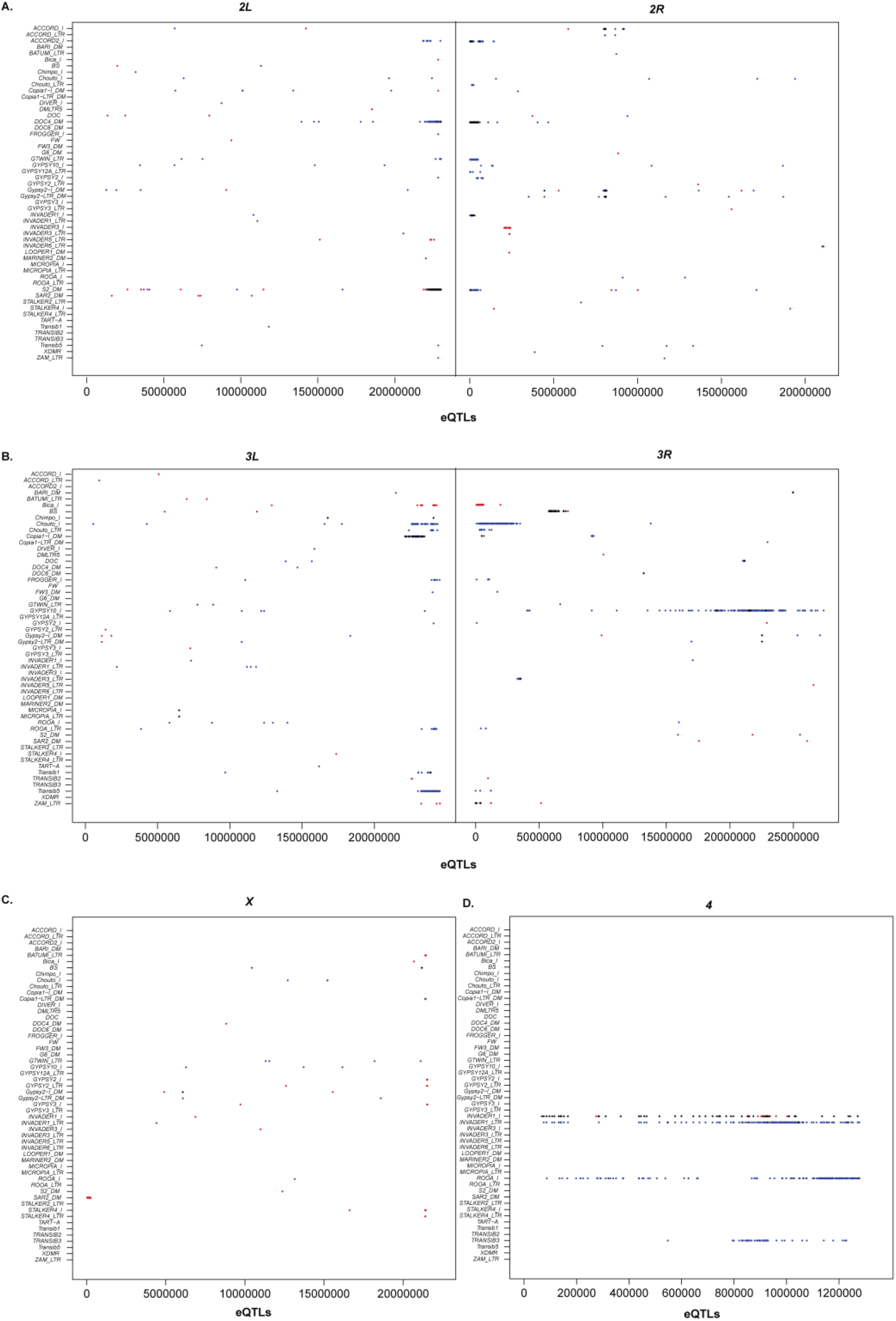
Genomic location of eQTLs for TE expression and associated TEs. eQTL chromosome positions (bp) are given on the *X*-axis, and the TEs with which they are associated on the *Y*-axis. Red points denote female-specific eQTLs, blue indicates male-specific eQTLs, and black shows eQTLs shared by males and females. (**A**) Chromosome *2*. (**B**) Chromosome *3*. **C**) Chromosome *X*. (**D**) Chromosome *4*.

**Figure S7.**
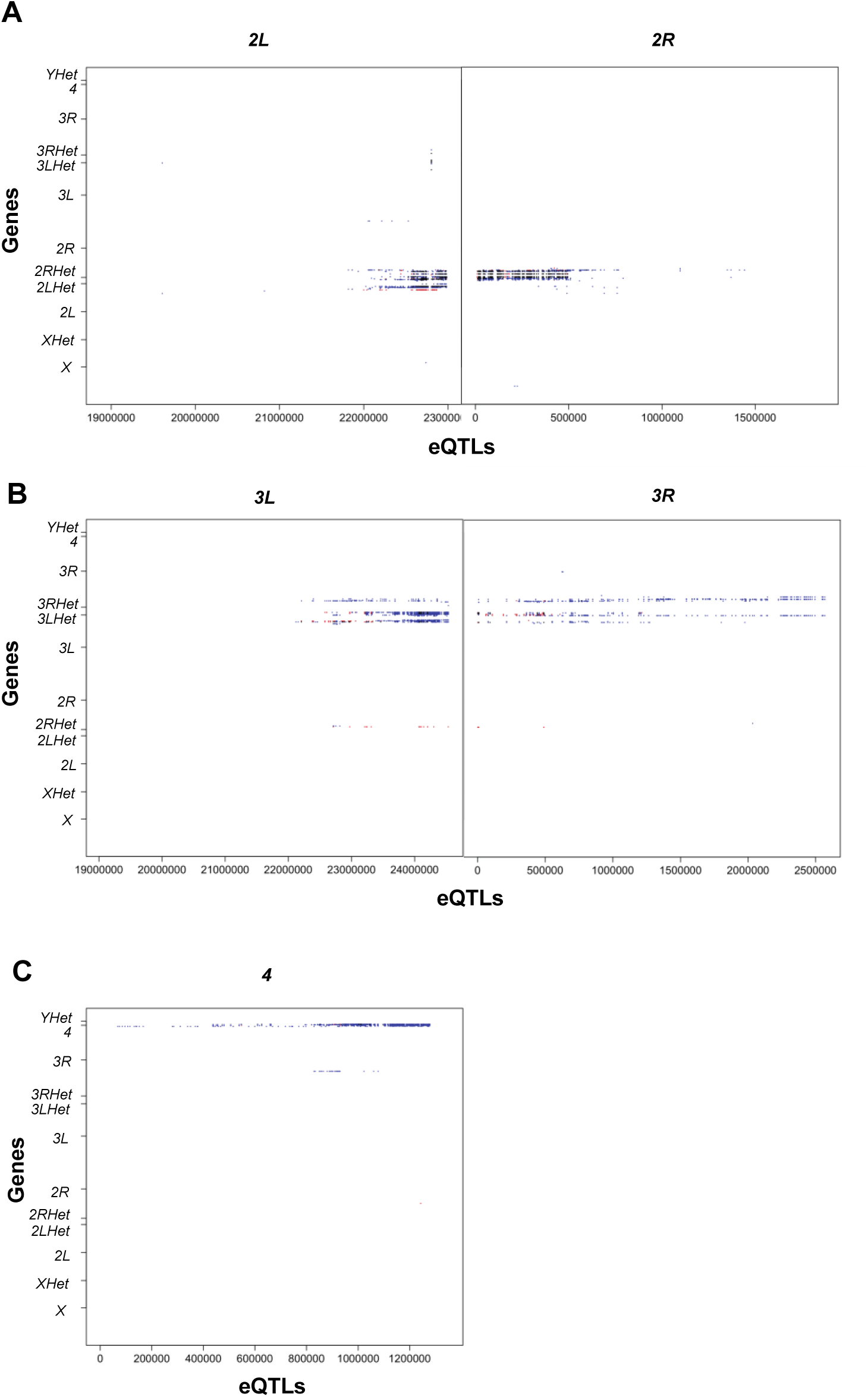
eQTL overlap between genes and TEs. eQTL positions are given on the *X*-axes, and the genes with which they are associated on the *Y*-axes. Red points denote female-specific eQTLs, blue indicates male-specific eQTLs, and black shows eQTLs shared by males and females. (**A**) Chromosome *2*. (**B**) Chromosome *3*. (**C**) Chromosome *4*.

## Supplementary Table Captions

**Table S1| Sequencing and alignment statistics for 800 RNA-seq samples.** Column legends are as follows. “Sample Name” format is DGRP Line Number, Sex (F = female, M = male) and Replicate (1 or 2). “Number of Sequencing Runs” denotes the number of sequencing runs in which the original sample library was sequenced in order to achieve sufficient sequencing depth. “Library Barcode” gives the Illumina barcode used for multiplex sequencing. “Total Reads Sequenced” gives the total reads in the raw fastq file generated by the Illumina Casava pipeline (after internal quality filtering). “Reads Removed by CutAdapt” and “% Reads Removed by CutAdapt” give the number and percent, respectively of reads removed by initial filtering with CutAdapt. “Reads Aligned to rRNA” and “% Reads Aligned to rRNA” give the number and percent, respectively, of reads identified as rRNA contamination by BWA. “Reads Aligned to Microbiome” and “% Reads Aligned to Microbiome” give the number and percent, respectively of reads aligned to the microbiome database by BWA. “Reads Aligned to RepBase” and “% Reads Aligned to RepBase” give the number and percent, respectively of reads aligned to RepBase by BWA. “Reads Aligned to *D. melanogaster* Genome” and “% Reads Aligned to *D. melanogaster* Genome” give the number and percent, respectively of reads uniquely aligned to the *D. melanogaster* reference genome by STAR.

**Table S2| Gene expression analyses.** (**A**) Mean expression (Log2 normalized FPKM values) for females and males across all DGRP lines for each known gene model from FlyBase. (**B**) Genomic coordinates, classification, and mean expression (Log2 normalized FPKM values) for females and males across all DGRP lines for each novel transcribed region (NTR). (**C**) Coding potential prediction of NTRs. (**D**) Results of pooled sex mixed-effect models run for all expressed gene profiles, including alignment bias estimates. (**E**) Results of female-only mixed-effect models for all expressed gene profiles, including alignment bias estimates. (**F**) Results of male-only mixed-effect models for all expressed gene profiles, including alignment bias estimates. (**G**) Chromosomal locations of genetically variable annotated genes (FBgn) and NTRs (XLOC).

**Table S3| Modules of genetically correlated gene expression.** WGCNA modules identified from within-sex line means of genetically variable gene expression levels, including the number of NTRs in each module; significantly enriched (5% FDR) Gene Ontology terms; Kegg and Reactome pathway membership; and Interpro protein domain annotation for known genes in each module, based on (**A**) female gene expression line means and (**B**) male gene expression line means.

**Table S4| Gene eQTL analyses.** (**A**) Female *cis*-eQTLs. (**B**) Male *cis*-eQTLs. Note that coordinates are given for Release 5 such that boundaries can be defined according to recombination map (see **C**). Release 6 coordinates are given in Table S2A. (**C**) Statistical tests for eQTL clustering, by chromosome. “Middle” denotes euchromatic regions with normal recombination and “edge” denotes euchromatic regions with reduced recombination according to Fiston-Lavier and Petrov‟s *Drosophila melanogaster* recombination rate calculator (http://petrov.stanford.edu/cgi-bin/recombination-rates_updateR5.plPetrov ref). (**D**). Numbers of eQTLs per gene (females). (**E**) Numbers of eQTLs per gene (males). (**F**) Statistical tests for enrichment for genes with more than 200 eQTLs and those with 199 or fewer eQTLs.

**Table S5| *cis-trans* eQTL networks.** (**A**) Female genes with *cis*- and *trans*-eQTLs. (**B**) Male genes with *cis*- and *trans*-eQTLs. (**C**) Female *cis-trans* eQTL networks. (**D**) Male *cis-trans* eQTL networks. (**E**) Overlap of genes in *cis-trans* eQTLs networks in males and females.

**Table S6| TE expression analyses.** (**A**) Log2 normalized RPM values for reads from each DGRP RNA-seq sample (columns) uniquely aligning to each known TE sequence in the *D. melanogaster* portion of RepBase. (**B**) Log2 normalized RPM values for reads from each DGRP line DNA-seq sample (columns) uniquely aligning to each known TE sequence. (**C**) Results of pooled sex mixed-effect models run for all TE sequences profiles in (**A**), including DNA copy number effects based on line profiles in (**B**), and copy number-independent line effects and line by sex interactions. (**D**) Results of female-only mixed-effect models for all TE sequences, including DNA copy number effects and copy number-independent line effects. (**E**) Results of male-only mixed-effect models for all TE sequences, including DNA copy number effects and copy number-independent line effects. (**F**) Female line means of copy-number independent effects inferred from the mixed-effect models in (**D**), for all TE sequences with significant LINE effects at 5% FDR threshold. (**G**) Male line means of copy-number independent effects inferred from the mixed-effect models in (**E**), for all TE sequences with significant LINE effects at 5% FDR threshold. (**H**) Modules of genetically correlated TE sequence expression, based on line means in (**F**) and (**G**), identified by WGCNA.

**Table S7| TE eQTL analyses.** (**A**) Female TE eQTLs. (**B**) Male TE eQTLs. (**C**) Summary of eQTLs by TE sequence. (**D**) Statistical tests by TE sequence for enrichment of eQTLs in euchromatic regions of normal recombination (“middle”) and pericentromeric euchromatin in which recombination is suppressed (“edge”), based on Fiston-Lavier and Petrov‟s *Drosophila melanogaster* recombination rate calculator (http://petrov.stanford.edu/cgi-bin/recombination-rates_updateR5.plPetrov ref). (**E**) Statistical tests by chromosome for enrichment of TE eQTLs in euchromatic regions of normal recombination (“middle”) and pericentromeric euchromatin in which recombination is suppressed (“edge”), based on Fiston-Lavier and Petrov‟s *Drosophila melanogaster* recombination rate calculator (http://petrov.stanford.edu/cgi-bin/recombination-rates_updateR5.plPetrov ref). (**F**) Female eQTLs associated with expression of multiple TE sequences. (**G**) Male eQTLs associated with expression of multiple TE sequences. (**H**) GO enrichment for genes with eQTLs associated with TE expression.

**Table S8| eQTLs associated with genes and TEs.** (**A**) Females. (**B**) Males.

**Table S9| Microbial species detected in DGRP RNA-seq libraries.** For each individual species, the genus is noted (where applicable), and the NCBI Taxonomic ID and Refseq genome assembly accession numbers are given for all genome assemblies included. Note that for some species there are multiple Taxonomic IDs (multiple known strains) and/or multiple genome assemblies available. The last column provides the total number of reads uniquely aligned to each microbial species summed across all DGRP RNA-seq samples, after removing all reads that align ambiguously to multiple microbial species or align to both microbial genomes and the assembled chromosomes of the *D. melanogaster* genome.

**Table S10| Microbial RNA expression analyses.** (**A**) Log2 normalized RPM (reads per million) values for reads from each DGRP RNA-seq sample (columns) uniquely aligning to each microbial species (rows). For *Aspergillus terreus* and *Malassezia globosa*, the majority of reads aligned to homologous regions of both species, and therefore these two species were combined for the purpose of this analysis. (**B**) Results of pooled sex mixed-effect models run for all species profiles in (**A**). For *Wolbachia pipientis*, a model was run without an additional factor for known Wolbachia infection status. For all other individual species, *P*-values were corrected for multiple testing using the Benjamini-Hochberg method and the corrected *P*-values are noted in corresponding FDR columns. (**C**) Line means, averaged across males and females, inferred from the mixed-effect models in (**B**), for all species with significant Line effects at a 5% FDR threshold. (**D**) Modules of genetically correlated microbial species, based on line means in (**C**), identified by WGCNA.

**Table S11| eQTLs for microbial species.** (**A**) eQTLs for microbial species (FDR ≤ 0.05). (**B**) eQTLs for microbial species (*P* < 10^−5^). (**C**) eQTLs associated with multiple microbial species (*P* < 10^−5^). (**D**) GO enrichment for genes with eQTLs associated (*P* < 10^−5^) with microbe expression.

**Table S12| Mean quantitative trait values for each DGRP line.** (**A**) Females. (**B**) Males.

**Table S13| ANOVA results for metabolic and body size traits.**

**Table S14| Most significant (P < 10^−5^) variants associated with quantitative traits from GWA analyses.** Variants highlighted in green are also eQTLs for gene expression. (**A**) Males. (**B**) Females.

**Table S15| Results of TWAS analyses.** Highlighted cells have transcript-trait associations with FDR ≤ 0.05. (**A**) Male genes (*P* < 10^−3^). (**B**) Female genes (*P* < 10^−3^). (**C**) Male TEs (*P* < 0.05). (**D**) Female TEs (*P* < 0.05). (**E**) Male microbial species (*P* < 0.05). (**F**) Female microbial species (*P* < 0.05).

