## Supplementary figures and images for "Gene Expression Networks in the Drosophila Genetic Reference Panel"

### Supplemental Figure 1

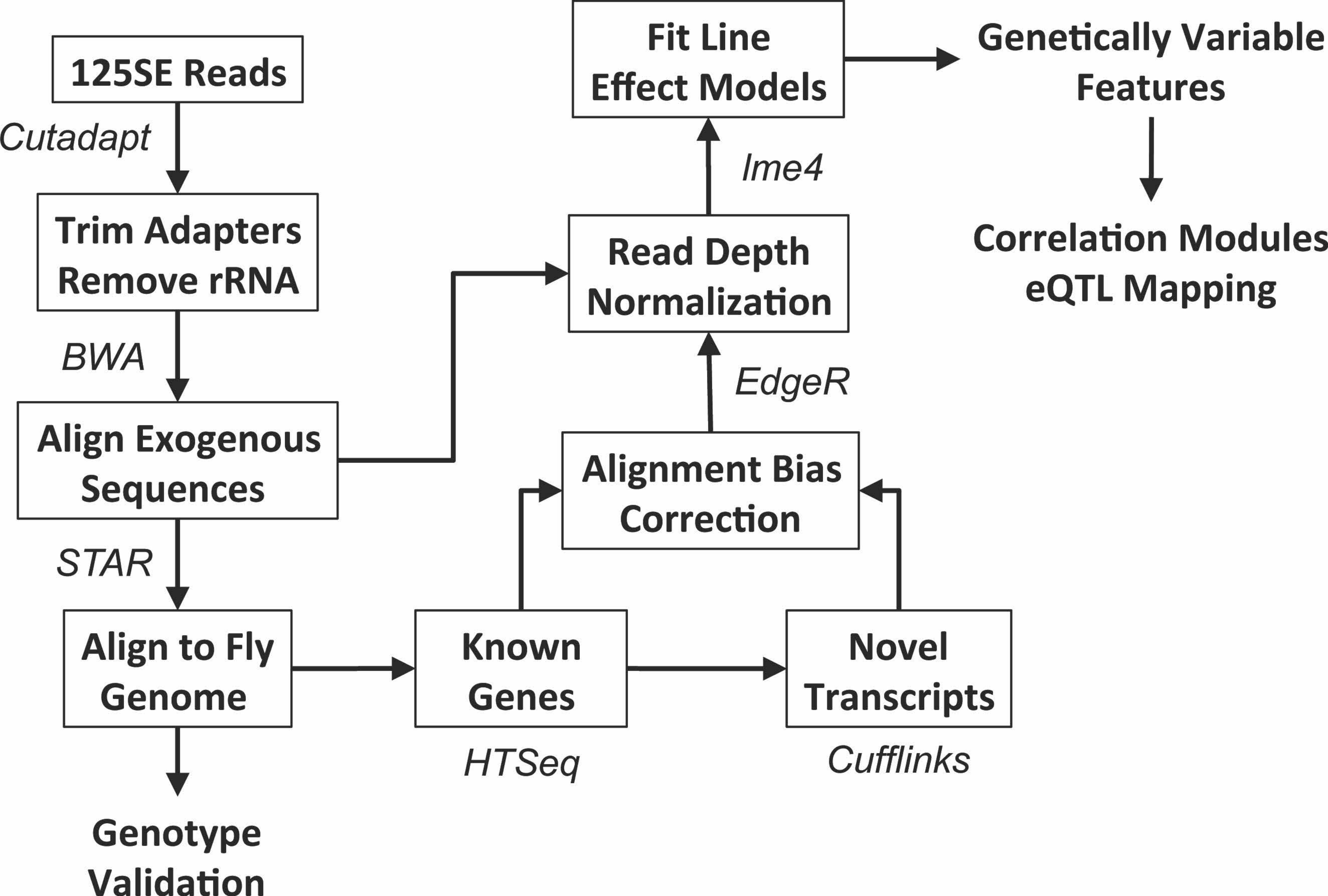

### Supplemental Figure 2

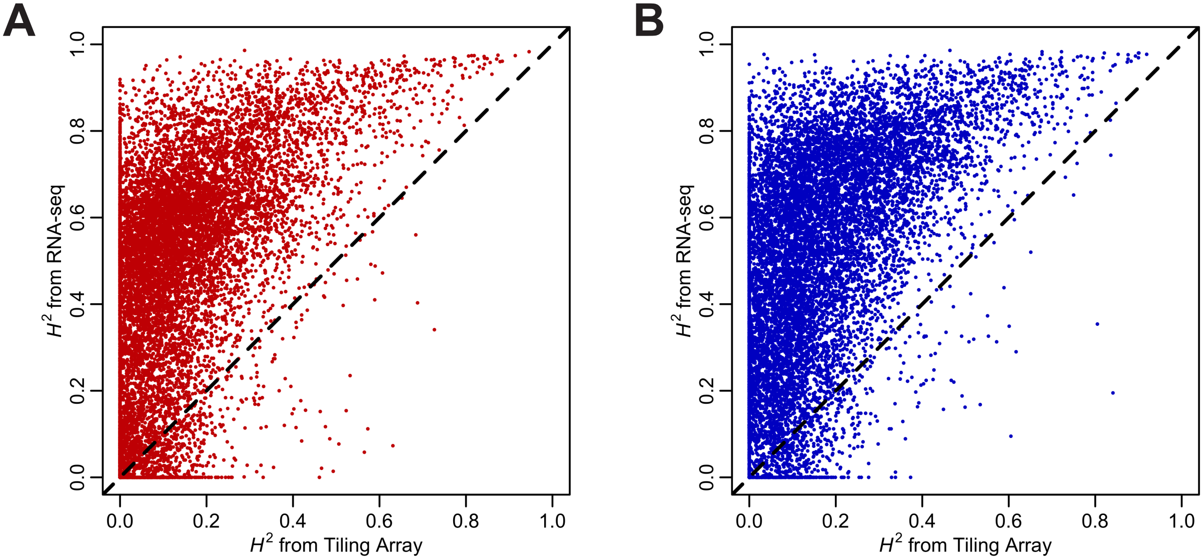

### Supplemental Figure 3

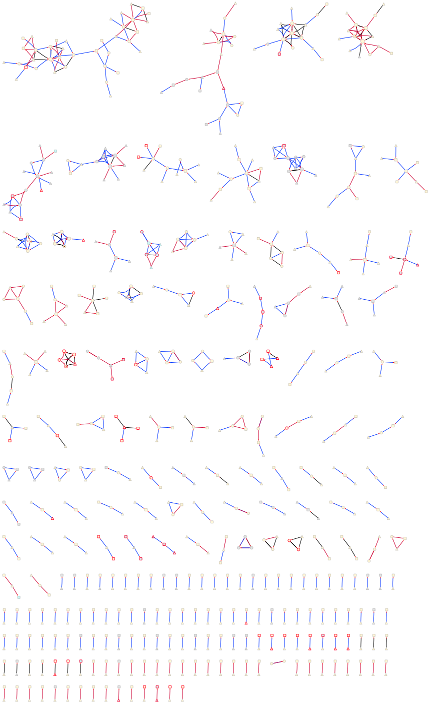

### Supplemental Figure 4

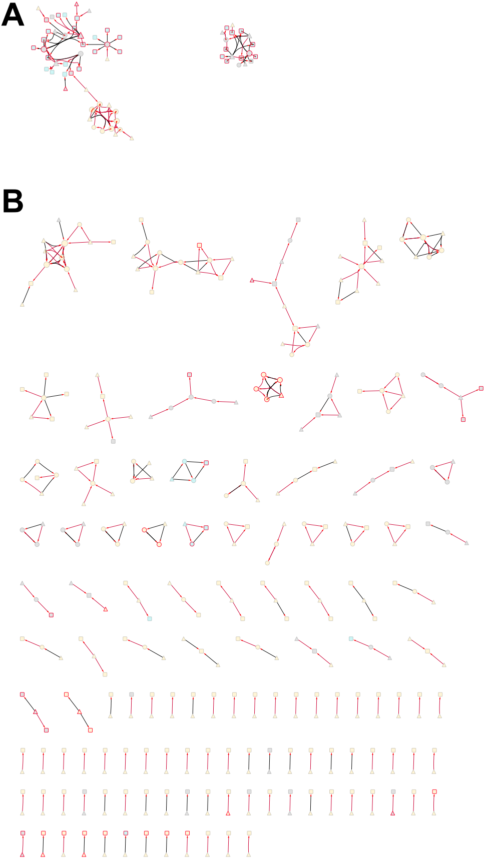

### Supplemental Figure 5

**A**

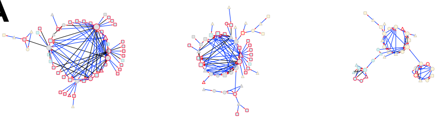

**B**

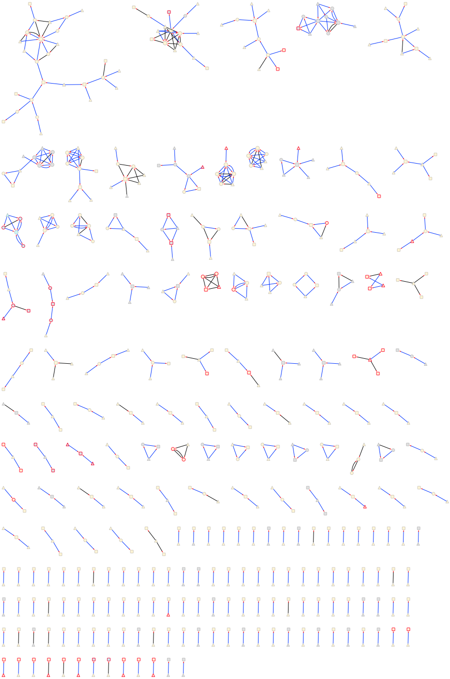

### Supplemental Figure 6

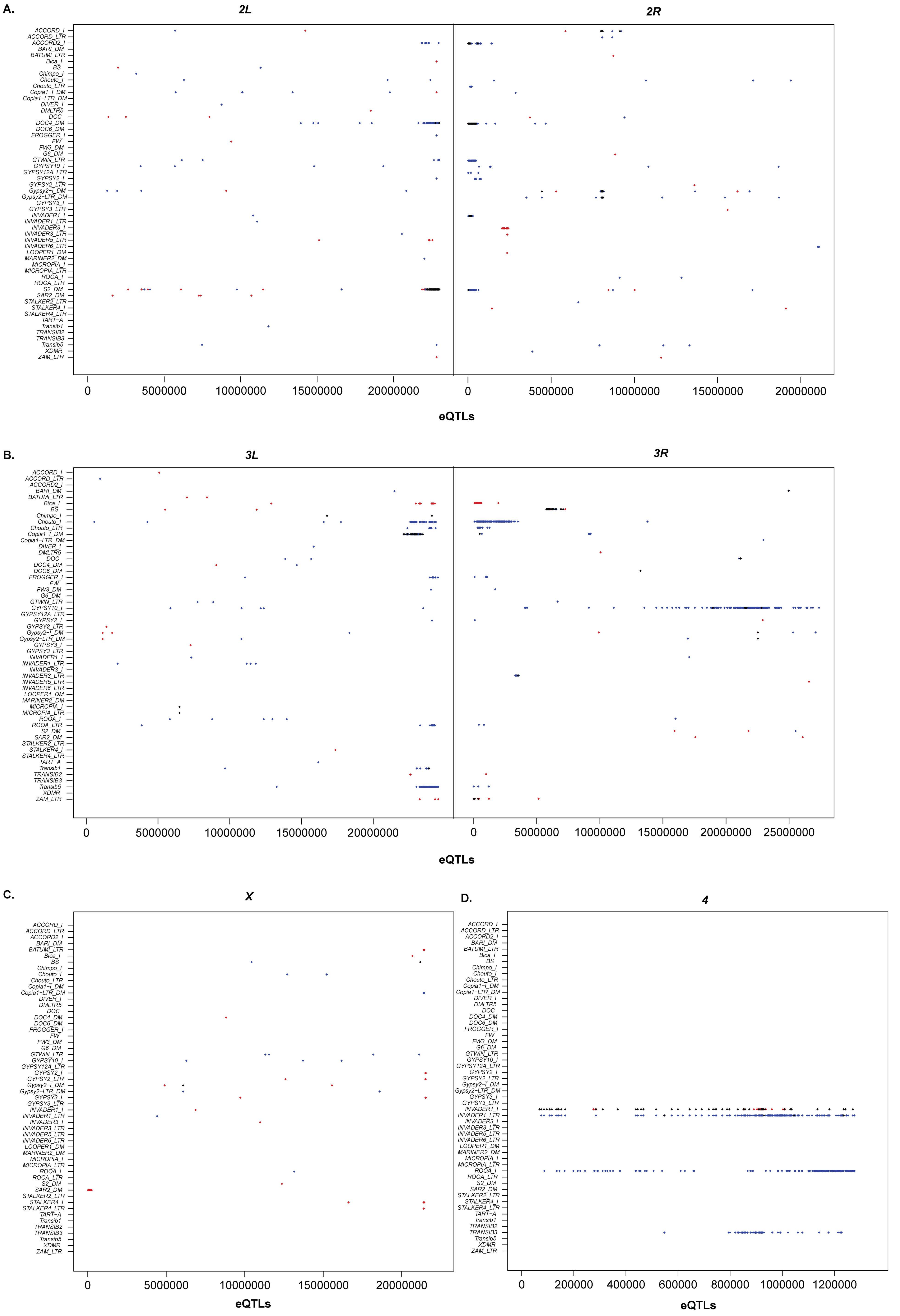

### Supplemental Figure 7

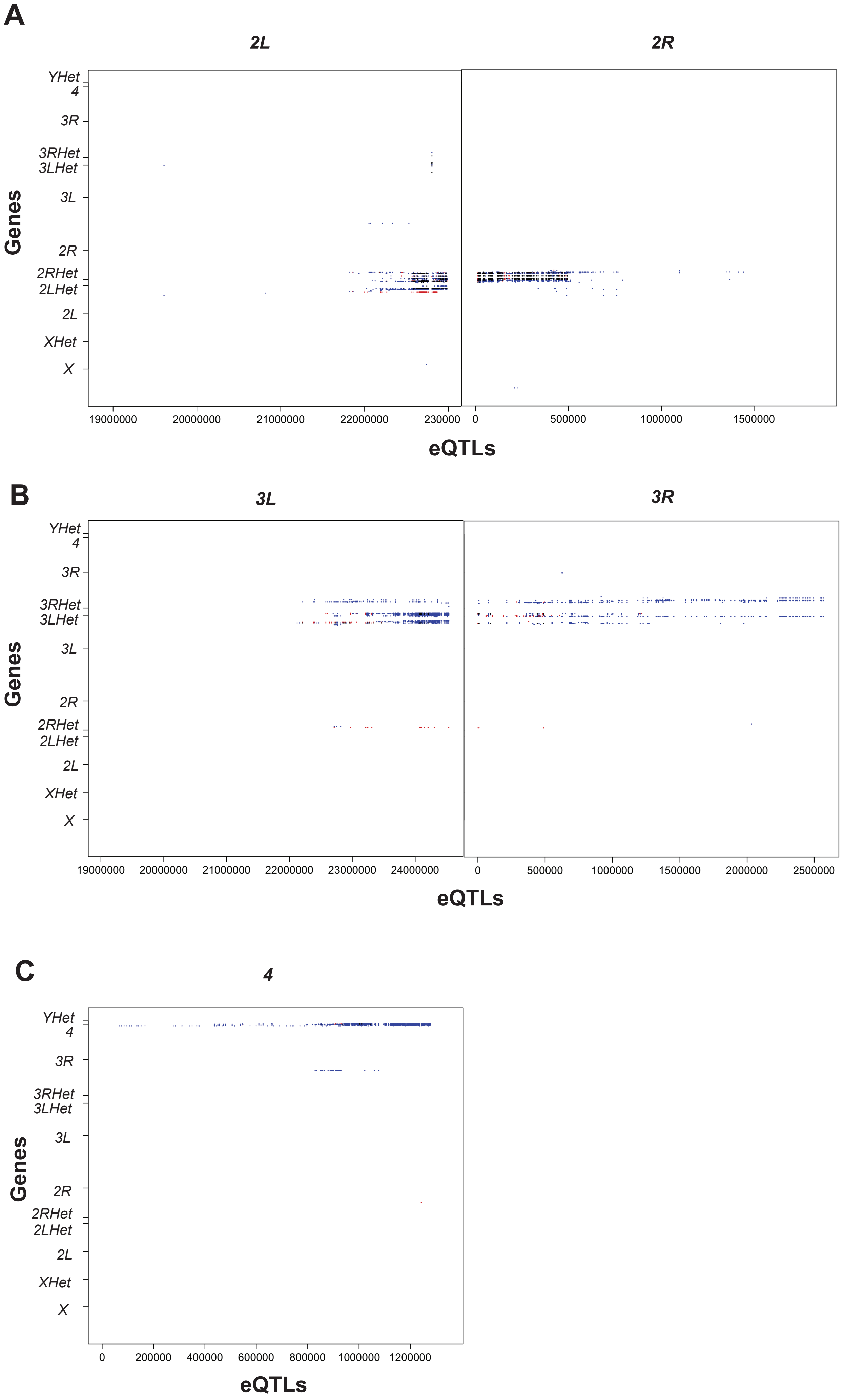
